# Specific cell states underlie complex tissue regeneration in spiny mice

**DOI:** 10.1101/2025.02.10.637521

**Authors:** Emilio Oviedo Rivadeneira, Robyn Allen, Mike Adam, Ashley W. Seifert

**Affiliations:** Department of Biology, University of Kentucky, Lexington, KY 40506, USA; Division of Experimental Hematology and Cancer Biology, Cancer and Blood Diseases Institute, Cincinnati Children’s Hospital Medical Center, Cincinnati, OH, USA; Department of Veterinary Anatomy and Physiology, University of Nairobi, Nairobi, Kenya; Aligning Science Across Parkinson’s (ASAP) Collaborative Research Network, Chevy Chase, MD 20815, USA

**Keywords:** Cell proliferation, regeneration, cell cycle, cell state, fibroblasts, chondroblasts

## Abstract

**SUMMARY STATEMENT:** Comparing regenerative vs. fibrotic healing, we identify injury-induced cell states associated with persistent cell cycle progression and complex tissue regeneration in mammals.

Cell proliferation is an elemental feature of epimorphic regeneration in vertebrate taxa. We previously reported that in contrast to fibrotic repair observed in laboratory mouse (*Mus*) strains, highly regenerative spiny mice (*Acomys* spp.) exhibit cell cycle progression and cell proliferation to faithfully replace missing tissue. However, little is known about proliferation dynamics, and specific cell types and states that may contribute to complex tissue regeneration in mammals. Using temporal pulse-chase experiments, we show that stromal cells in *Acomys dimidiatus* rapidly re-enter the cell cycle in response to injury and maintain tight spatiotemporal control of cell cycle progression to restrict the proliferative population to a distal area relative to the injury. Conversely, *Mus* stromal cells incorporate thymidine analogs without cell division supporting an S-phase arrest after D10. Deploying immunostaining and scRNA-seq, we identify several key cell types (CRABP1+, αSMA+) differentially associated with regenerating versus scar tissue. Importantly, our single cell data revealed distinct gene expression profiles for cross-species stromal cell types, identifying cell states specific for regenerative or fibrotic healing. While CRABP1+ fibroblasts are enriched in *Acomys* ears before and after injury, similar fibroblasts enriched in young, postnatal *Mus* ears remain unable to promote regeneration. Our data underscore the finely regulated dynamics of proliferating cells during regeneration and emphasize that regeneration depends on multiple factors including the presence of specific cell types and the ability of cells to acquire specific states.

**Key Conclusions:**

-Differentiated cells in *Acomys*, *Mus* and *Danio* re-enter the cell cycle in response to injury, while homeostatic cycling cells contribute to blastema formation in *Ambystoma*
-Pulse-chase thymidine analog labeling shows tight spatiotemporal control of proliferating stromal cells during regeneration in *Acomys*.
-Following injury, CRABP1 and αSMA are expressed in distinct stromal cell populations in *Acomys* but are co-expressed in *Mus* stromal cell populations.
-Species-specific cell states underlie regenerative and fibrotic repair
-CRABP1+ cells are lost during embryonic development in *Mus* ear pinna but are retained in *Acomys* to adulthood.
-Young neonatal *Mus* with abundant CRABP1+ cells still fail to execute regenerative healing

## INTRODUCTION

Epimorphic regeneration is mediated by proliferating cells that form a blastema and subsequently differentiate to replace missing tissue (Morgan, 1901). Nearly all regenerative species use epimorphic regeneration to rebuild missing tissue. Seminal studies using urodeles, zebrafish, and planarians explored the dynamics of proliferating cells to understand their origin, proliferative rates, migration patterns, and responses to different injury types (Hays & Fischman, 1961; Poleo et al., 2001; Wallace, 1981; Wenemoser & Reddien, 2010). For example, thymidine analogs applied to amputated newt limbs revealed that cells start to synthesize DNA in differentiated tissues adjacent to the wound site (Hays & Fischman, 1961). These cells are then liberated and displaced distally to form the blastema. Similar studies in axolotls quantified cell cycle length, identified factors that maintain proliferation, and blocked proliferation to study regeneration (Maden & Wallace, 1976; Wallace, 1981; Wallace & Maden, 1976). In planarians, different injury types trigger distinct mitotic responses. For example, amputation causes local and systemic responses, while injuries that do not remove tissue only trigger a systemic response (Wenemoser & Reddien, 2010). Similarly, zebrafish regeneration studies have characterized the spatiotemporal distribution of proliferating cells across the blastema (Nechiporuk & Keating, 2002; Poleo et al., 2001). Considering these studies, an open question is the degree to which patterns and processes governing cell proliferation may hold during instances of epimorphic regeneration in mammals.

Complex tissue regeneration in vertebrates has been primarily studied in fishes, amphibians and invertebrates. However, some mammals exhibit bona fide regeneration (Billingham et al., 1959; Borgens, 1982; Carlson, 2005; Gawriluk et al., 2016; Goss & Grimes, 1972; Neeham, 1953; Seifert et al., 2012; Vorontsova et al., 1960). For instance, patterned musculoskeletal tissue is reconstituted after removal with a full thickness hole punch through the ear pinna in some lagomorphs and deomyine rodents (e.g., *Acomys* and *Lophuromys*) (Gawriluk et al., 2016; Goss, 1980; Goss & Grimes, 1972; Joseph & Dyson, 1966; Riddell et al., 2025; Seifert et al., 2012). Comparative studies between regenerative and non-regenerative rodents showed that spiny mice (*Acomys*) can regenerate ear pinna tissue after full thickness excisional injury; a response mediated by cell proliferation and blastema formation. In response to an identical injury, however, laboratory mice (*Mus*) cannot regenerate missing tissue and while cells are activated to re-enter the cell cycle in response to injury, these cells are arrested shortly afterwards (Gawriluk et al., 2016; Seifert et al., 2012).

In adult mammals, cell division is generally restricted to stem and progenitor cells in tissues that require constant turnover during homeostasis (Cheung & Rando, 2013; Liu et al., 2019; Orford & Scadden, 2008). In highly regenerative mammals, are there certain cell types or states that can specifically re-enter the cell cycle and proliferate? In highly regenerative vertebrates, blastema formation appears to be supported by cellular re-programming (aka dedifferentiation) of lineage-restricted progenitors located near the wound site (Echeverri et al., 2001; Gerber et al., 2018; Jopling et al., 2010; Leigh et al., 2018). Interestingly, while lineage-restricted progenitors undoubtably support mouse digit tip regeneration (G. L. Johnson et al., 2020; Lehoczky et al., 2011; Rinkevich et al., 2011), the capacity of mesenchymal cells to maintain a specific transcriptomic state is also associated with regeneration (Storer et al., 2020). Similar observations come from research comparing injured deer antler velvet (regenerative) to dorsal back skin (non-regenerative) where both skin types have *Acta2*+ myofibroblast cells, but specific transcriptomic profiles in these cells are associated with location. Importantly, velvet fibroblasts access regulatory networks skewing them towards an immunosuppressive, pro-regenerative fetal phenotype, while back skin fibroblasts express networks related to an inflammatory, pro-fibrotic outcome (Sinha et al., 2022). Wound healing studies in large skin wounds that regenerate hair follicles are biased to use a specific fibroblast subtype that is inhibited in fibrotic responses; a finding that is also associated with better healing outcomes in neonatal *Mus* and human skin (Guerrero-Juarez et al., 2019; Mine et al., 2008; Phan et al., 2021a). Together, these studies support that access to regenerative cell states ***and*** the presence of specific cell types is essential to promote a regenerative outcome.

Here we use the spiny mouse ear pinna regeneration model compared to fibrotic repair of identical injuries in laboratory mice to explore the spatiotemporal dynamics of proliferating cells and begin identifying cell types that participate in regeneration. First, we use a pulse-chase method to quantify cell cycle activation in regenerative vertebrates with determinate and indeterminate growth and find that growth mode is not strictly associated with regenerative ability or the lack thereof. Additionally, we find no evidence of long-range cell activation in response to injury. Using a pulse-chase labeling strategy in *Acomys*, we show that proliferating cells maintain a distal position relative to the wound site, while cells in the proximal compartment exit the cell cycle and differentiate. Finally, using a combination of immunostaining and scRNA-sequencing, we identify similar cell types across species that express distinct transcriptomic states associated with regenerative or fibrotic repair supporting that species-specific cell states underlie regeneration and scaring.

## METHODS

### Animal care

Sexually mature, spiny mice (*Acomys dimidiatus* - aged six months to one year old) and *Mus musculus*, ∼three months old) were obtained from our in-house breeding colonies at the University of Kentucky. Animals were maintained on a 12:12 h L:D light cycle. Both species were fed 14% mouse chow (Teklad Global 2014, Harlan Laboratories) and *Acomys* diets were supplemented with black sunflower seeds. All animal experiments were approved by University of Kentucky Institutional Animal Care and Use Committee (IACUC) under protocol 2013–3254.

### 4mm ear punch assay regeneration assay

A 4mm full thickness excisional punch through the ear pinna serves as a tissue regeneration assay (Gawriluk et al., 2016). Animals were anesthetized with 3% vaporized isoflurane (v/v) (Henry Schein Animal Health, Dublin, OH) at 1 psi oxygen flow rate. A 4mm biopsy punch (Sklar Instruments, West Chester, PA) was used to create a through-and-through hole in the right and left ear pinna. Ear tissue was collected at specified time points using an 8mm biopsy punch (Sklar Instruments, West Chester, PA) circumscribing the original injury. The published protocol can be found here: dx.doi.org/10.17504/protocols.io.bp2l6dx65vqe/v1.

### Pulse-chase experiments

We injected Ethynyl deoxyuridine (EdU) (10 ug/g) and Bromodeoxyuridine (BrdU) (1 ug/g) diluted in normal saline, for pulse-chase experiments to label and track cycling cells (the published protocol can be found here: dx.doi.org/10.17504/protocols.io.261gerdmol47/v1). Both solutions were injected intraperitoneally with a difference of five days between the EdU and BrdU injections. EdU and BrdU injections were given as three separate injections, one every three hours over a period of nine hours. The left ear pinna was collected after the EdU pulse to assess baseline cell numbers in S-phase, and the right pinna was collected after the BrdU pulse to track single (EdU+ or BrdU+) and double positive cells. The injection scheme was as follows: the first group of animals was injected with EdU at D0 and BrdU at D5, the next group was injected with EdU at D5 and BrdU at D10 and so on until D30. Hence, tissue samples were collected at 0, 5, 10, 15, 20, 25, and 30-days post injury.

Our experimental scheme allowed us to distinguish cells that (i) were in S-phase at the time of the EdU injections and stopped proliferating within next five days (Edu+ alone); (ii) were in S-phase at the time of the EdU injections, divided multiple times over five days diluting the EdU, and remained in the cell cycle acquiring BrdU (weak EdU+/ strong BrdU+) - because each cell division dilutes the EdU by half, a scenario where EdU is not detectable is possible, and these cells would contribute to the BrdU+ alone counts); (iii) cells not in S-phase during the EdU pulse but were cycling during the BrdU injections (BrdU+ alone); (iv) cells that acquired EdU, divided only a few times, and remained in the cell cycle during the BrdU chase (strong EdU+/ strong Brdu+).

### Histology, EdU detection and Immunohistochemistry

Harvested tissue was fixed overnight in 10% neutral buffered formalin (NBF) (Thermo Fisher Scientific Inc. Epredia 5701) at 4°C on a shaker. After 16-20 hrs the tissue was washed 3x with PBS, 3 minutes per wash, followed by 3x 70% ethanol washes for dehydration and storage at 4°C. Stored tissue was processed for paraffin (Leica Biosystems, Buffalo Grove, IL, Formula “R” REF: 3801450) embedding using a Histo5 tissue processor machine (Milestone). Samples were sectioned at 5μm. EdU was detected using click chemistry as follows: Tissue sections were de-paraffinized and rehydrated. Antigen retrieval was performed if co-stained with antibodies. Tissues were washed two times for 10 minutes in Tris-buffered saline (TBS) to incubate in the prepared reaction cocktail (see table below) for 30 minutes at RT protected from light. Tissues were washed two times in TBS for 5 minutes and if co-stained with antibodies were blocked following the fluorescent IHC protocol for BrdU and cell type specific antibodies tissue sections were deparaffinized and re-hydrated. Tissues were then exposed to high heat antigen retrieval in a steamer in antigen unmasking TRIS or citric acid-based solutions (Vector Laboratories, Inc.). After a 5-minute wash, tissues were blocked in blocking serum (see table below) for 30 minutes at RT. Tissues were then incubated with primary antibody diluted in blocking serum at 4 °C overnight. After a 5 minutes wash, tissues were incubated in secondary antibody diluted in blocking serum for 30 min at RT. Tissue samples were incubated with Hoechst 33342 for 5 minutes at RT and then dried and mounted in Prolong. Finally, protein similarity was compared to assess antibody binding efficacy (Supplementary table S1). For complete protocol details regarding paraffin sections (dx.doi.org/10.17504/protocols.io.6qpvr9353vmk/v1), immunohistochemistry (dx.doi.org/10.17504/protocols.io.n92ldrmd8g5b/v1), and EdU detection on sections (dx.doi.org/10.17504/protocols.io.e6nvwbdrzvmk/v1).

| **EdU reaction cocktail** |  |
| --- | --- |
| **Components** | **Volume (ul)** |
| 2 M Tris, pH 8.5 | 5 |
| 50 mM CuSO<sub>4</sub> | 2 |
| 0.5 M ascorbic acid | 20 |
| Alexa fluor azide (0.25 mg/ml) | 0.2 |
| ddH<sub>2</sub>O | 73 |
| Total | 100 |

| **Primary Antibodies** |  |  |  |  |  |
| --- | --- | --- | --- | --- | --- |
| **Antibody** | **Company, Cat#** | **RRID** | **Host species** | **Dilution** | **Antigen retrieval** |
| Anti-BrdU (Bromodeoxyuridine) | Abcam, Ab6326 | AB_305426 | Rat | 1:500 | Citric Acid |
| Anti-wide spectrum Cytokeratin | Abcam, Ab9377 | AB_307222 | Rabbit | 1:500 | None |
| Anti-ionized calcium-binding adapter molecule 1 (IBA1) | Novus Biologicals, NB100-10288s | AB_521594 | Goat | 1:400 | Tris |
| Anti-retinoic acid-binding protein 1 (CRABP1) | Cell Signaling Technology, Inc, 13163 | AB_2750569 | Rabbit | 1:500 | Tris |
| Anti- alpha-Smooth Muscle Actin (alpha-SMA) | Novus Biologicals, NB300-978 | AB_2273628 | Goat | 1:200 | Tris |
| Anti-SRY Box Transcription Factor 9 (SOX9) | Abcam, ab185230 | AB_2715497 | Rabbit | 1:1000 | None |
| Anti-macrophage mannose receptor (MMR) (CD206) | R&D Systems, AF2535 | AB_2063012 | Goat | 1:500 | Tris |

| Secondary Antibodies |  |  |  |  |
| --- | --- | --- | --- | --- |
| Antibody | Company, Cat# | RRID | Host species | Dilution |
| Alexafluor 594 anti-Rabbit | Thermo Fisher Scientific, A21207 | AB_141637 | Donkey | 1:400 |
| Alexafluor 647 anti-goat | Thermo Fisher Scientific, A21447 | AB_2535864 | Donkey | 1:400 |
| Alexafluor 647 anti-rat | Thermo Fisher Scientific, A21247 | AB_141778 | Goat | 1:400 |

### Western Blotting

Protein extraction and quantification was performed as indicated (dx.doi.org/10.17504/protocols.io.rm7vzkx12vx1/v1). Western blotting was performed as indicated (dx.doi.org/10.17504/protocols.io.kqdg3qxz1v25/v1).

### Imaging and Cell Quantification

Fluorescent images were collected using Cells Sens from Olympus in Olympus inverted microscope (Olympus America Inc). Whole tissue sections were captured in 20X and stitched automatically. Cell counting was manually performed in defined areas of the images separately in each channel using ImageJ. Percentages of positive cells were calculated by quantifying nuclei (Hoechst 33342) using the threshold tool in ImageJ.

### Statistics

Statistical analyses were performed using JMP (version Pro 15.0, SAS Institute Inc) and SPSS (version 28.0, IBM) where alpha was set at 0.05 for all statistical analyses. Graphs were made in excel and JMP. To analyze our pulse chase data across species, we performed Two-way ANOVAs with either EdU+, BrdU+ or double positive cells (dependent variable), and days post injury (independent variable) followed by Tukey HSD multiple comparisons to test for differences in positive cells as a function of time (Supplementary table S2).

To analyze proliferation during regeneration in *Acomys*, we used a linear mixed model to assess differences in percentage of positive cells between three positions (A, B, C) over time (D5-D30) (Supplementary table S3). Likelihood ratio testing and Akaike Information Criterion (AIC) were used to select an appropriate covariance structure. A Kenward-Roger adjustment was used to correct for negative bias in the 170 standard errors and degrees of freedom calculations induced by the small sample size. Relevant pairwise comparisons were performed and controlled for multiple comparisons using Fisher’s Least Significant Differences (LSD). Mixed modeling analysis was performed using SAS 9.4 (SAS Institute, Inc.; Cary, NC, USA).

### Single-cell analysis

scRNA-seq data used in this study were download and available through the Gene Expression Omnibus (GEO) (RRID:SCR_005012) under the accession number GSE182141. The raw fastqs were processed through the CellRanger pipeline v8.0.1 (RRID:SCR_017344) using 10X Genomics’ mm10-2020-A reference genome or a custom annotated *Acomys dimidiatus* genome (GCA_907164435.1 (mAcoDim1_REL_1905), with the ‘include introns’ set to true to obtain gene expression matrices.

Background reads were removed using decontX from the celda package using the filtered barcodes as the cells to keep and the remaining cells in the raw barcode matrix as the background (Campbell et al., 2024). Doublets were minimized using DoubletFinder (RRID:SCR_018771) (McGinnis et al., 2019). The R v4.4 (RRID:SCR_001905) library Seurat v5.1 (RRID:SCR_016341) (Hao et al., 2021; Hao et al., 2024)) was used for cell type clustering and marker gene identification. Cells expressing >500 genes were retained for downstream analysis. Each sample was normalized SCTransform v2 (RRID:SCR_022146), using the glmGamPoi method and the number of RNA molecules per cell were regressed out. Samples were integrated with common anchor genes using the CCA method to minimize sample to sample variation. Cell clusters were determined by the Louvain algorithm using a resolution of 0.3. UMAP dimension reduction was done using the first thirty principal components. Marker genes for each cell type were calculated using the Wilcoxon Rank Sum test returning only genes that are present in a minimum of 25% of the analyzed cluster. Cross species markers were done as above in cell types of interest, but the pct.2 score was filtered to be > 0 to eliminate uniquely annotated genes from dominating the lists. Volcano plots were created with the Enhanced Volcano (Blighe et al., 2024) R library using the avg_log2FC and p_val_adj values as inputs.

Gene ontology (GO) analyses were performed in the Enrichr platform (https://maayanlab.cloud/Enrichr/). Only differentially expressed genes with a two-fold change were included in the analysis of GO Biological Process.

## RESULTS

### Vertebrate stromal cells rapidly re-enter the cell cycle in response to injury

Notably few regeneration papers using salamanders or fish have indicated whether stromal cells are proliferating during normal homeostasis. To explore proliferation in uninjured tissue, we used EdU to label cells in S-phase in two highly regenerative amniotes (*Danio rerio* - zebrafish, *Ambystoma mexicanum* - axolotl), in an adult mammal with enhanced regenerative ability (*Acomys dimidiatus* – spiny mouse) and in a poorly regenerating mammal (*Mus musculus* – outbred ND4 laboratory mouse strain) (Fig. 1A). Examining the ear pinna (*Acomys* and *Mus*), limb (*Ambystoma*) and caudal fin (*Danio*), we found cycling cells in the epidermis from all four species prior to injury; a finding that was expected given the presence of slow cycling stem cells in the epidermis (Fig. 1B). In uninjured *Danio* and *Ambystoma* tissue, we also found a small percentage of cycling cells in the stroma (1% and 0.5% respectively) supporting previous reports for these species (Fig. 1B-C) (K. Johnson et al., 2018; Simkin & Seifert, 2018). In contrast, we found only one or two cycling stromal cells across entire tissue sections in *Acomys* or *Mus* prior to tissue removal (Fig. 1B-C). These data support that stromal cell proliferation is essentially non-existent prior to injury in mammals, at least in the ear and skin.

**Figure 1.**
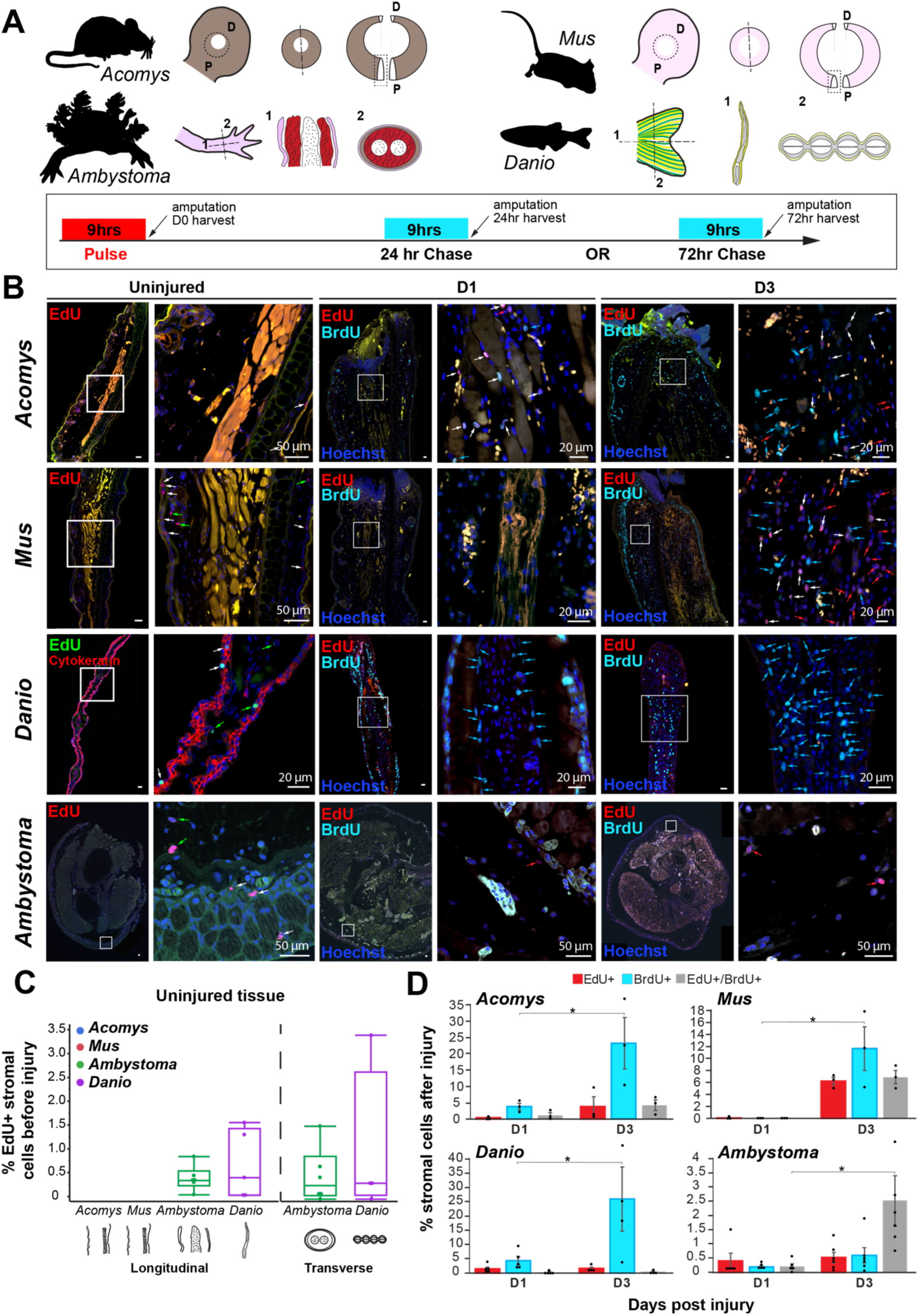
Cell cycle re-entry before and after injury in *Acomys*, *Mus*, *Danio* and *Ambystoma*. A) Tissue section planes and EdU/BrdU pulse-chase scheme indicating injections and harvest times; D = distal, P = proximal. B) Representative images of EdU+ and BrdU+ cells in uninjured tissue (D0), D1 and D3. In uninjured tissue, white arrows = positive cells in the epidermis and green arrows = positive cells in the stromal compartment. In injured tissue, red arrows = EdU+ cells, light blue arrows = BrdU+ cells, and white arrows = double positive cells (EdU+/BrdU+). C) EdU+ cell quantification as a percentage of the total number of cells in uninjured tissue from all four species. Autofluorescent tissue, including red blood cells, stains in multiple channels (yellow). D) EdU+, Brdu+ and EdU+/BrdU+ cell quantification as a percentage of the total number of cells at D1 and D3 post-injury. Images represent n = 3 biological samples/time point in *Acomys* and *Mus*. See Methods for cell counting quantification. In *Danio* (zebrafish), n = 9 at D0, n = 5 at D1 and n = 4 at D3. In *Ambystoma* (axolotl), n = 12 at D0, n = 6 at D1, n = 6 at D3. (* *p*<0.05, ** *p*<0.01).

A longstanding hypothesis is that cells in mammals are refractory to cell cycle re-entry and progression whereas cells in highly regenerative vertebrates easily re-enter the cell cycle and proliferate to replace missing tissue (Pajcini et al., 2010; Simkin & Seifert, 2018; Tanaka et al., 1997; Tanaka & Brockes, 1998). To test this hypothesis, we used a pulse chase strategy whereby we injected animals with EdU, performed the specified injury, injected BrdU at either 15 or 63hrs post injury (hpi) and then harvested tissue 9hrs later at 24 and 72 hpi (see Methods). We quantified stromal cells that were EdU+, or BrdU+, or EdU+/BrdU+ (double positive) (Fig. 1B, D). 24 hpi spiny mice and zebrafish showed the highest percentage of newly cycling cells in response to injury (BrdU+). *Mus* cells also re-entered the cell cycle, albeit later at 72 hpi. Axolotl limb tissue, on the other hand, exhibited very few singly BrdU+ cells but presented the highest number of double positive cells (Fig. 1D). These data demonstrate that stromal cells in *Acomys*, *Mus* and zebrafish re-enter the cell cycle *de novo* in response to injury with a 48hr lag observed for *Mus* cells.

To assess cell activation arising from the injury but independent of the local repair response (i.e., systemic cell activation), we collected the contralateral ear pinna in *Mus* and *Acomys*, the contralateral limb in axolotls, and the pelvic fin in zebrafish 24 hpi and 72 hpi after tissue amputation. Using a pulse chase strategy as described above, we looked for cell cycle re-entry in uninjured tissues at 24 hpi and 72 hpi. EdU+ cells were cycling prior to injury, BrdU+ cells were those activated systemically and EdU+/BrdU+ were those cycling prior to injury that were activated systemically. Despite reports in lab mice of long-range cell activation in skeletal muscle stem cells (Rodgers et al., 2014), in *Acomys* and *Mus*, we did not find a noticeable response in the contralateral ear pinna where the number of BrdU+ cells did not exceed 0.5% at either time point (Fig. S1A-B). Similarly, in axolotl and zebrafish, the number of EdU+ cells at 24 and 72 hpi was similar to the number of positive cells found in uninjured tissue at D0 (see above) (Fig. S1A-B). These data indicate that a systemic response to injury is negligible at 24 and 72hpi in all four species.

### Proliferating cells are maintained in a distal position relative to the wound site over time

*Acomys* species regenerate ear pinna tissue after removal with a 4 mm (or larger) biopsy punch whereas identical tissue excisions in wild or laboratory *Mus* result in scar tissue formation (Seifert et al., 2012). To more thoroughly understand the kinetics of proliferating cells during regeneration and fibrotic repair, we again employed a pulse (EdU) chase (BrdU) scheme where our pulse and chase were separated by a five-day window (see Methods). First, we analyzed BrdU+ cells to estimate the proliferative rate from injury until D30 and then analyzed the spatial distribution of cycling cells by quantifying cells adjacent to the injury (aka proximal) and at two distal locations (i.e., closest to the new epidermis); distal dorsal and distal ventral (Fig. 2A-E).

**Figure 2.**
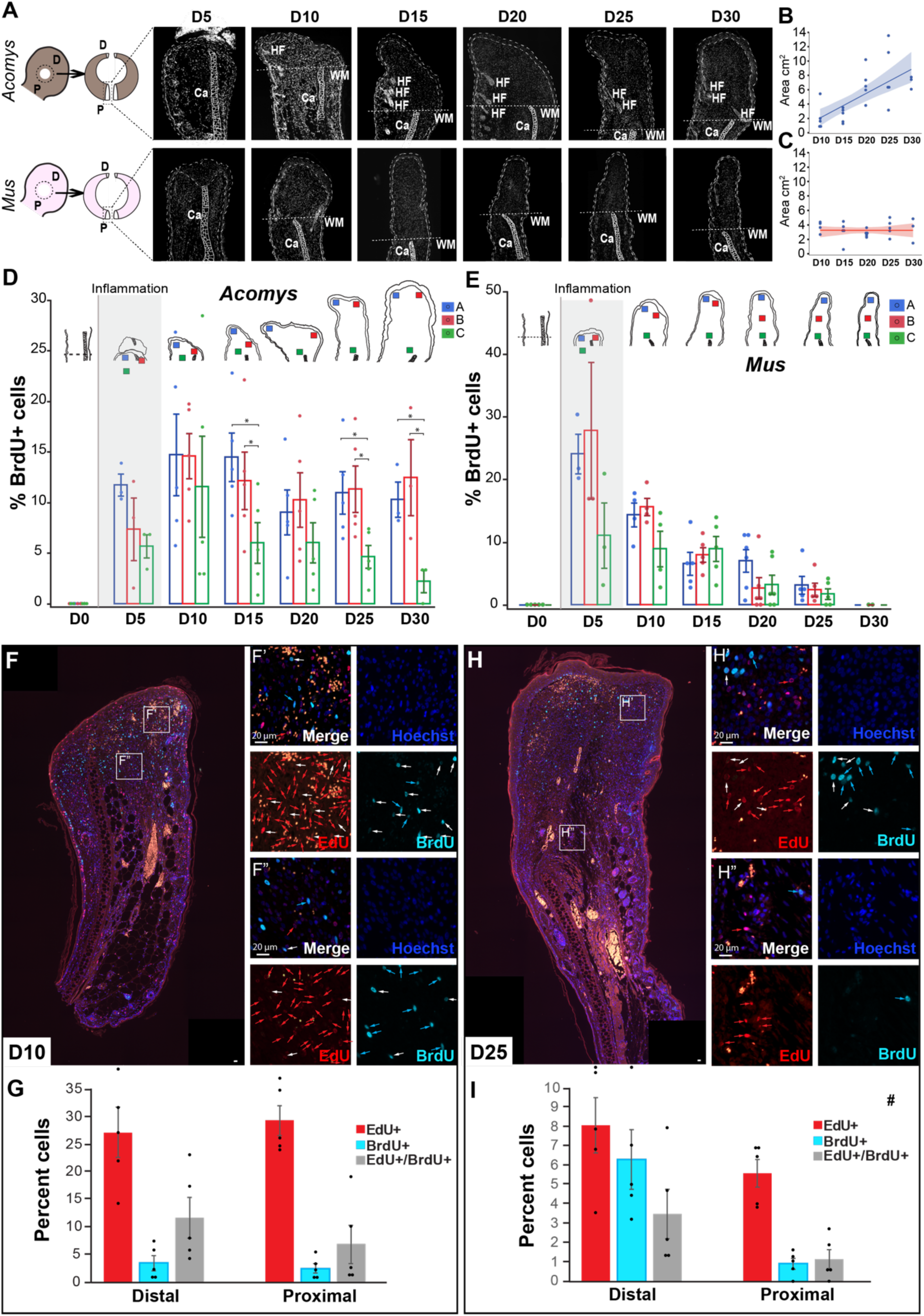
Spatiotemporal distribution of proliferating cells in *Acomys* and *Mus*. A) Representative images of blastema and scar tissue changes over time (D5-D30), Ca = cartilage, WM = wound margin, HF = hair follicle. B-C) Blastema or scar tissue area in mm^2^ from D10 to D30 based on tissue sections. D-E) Quantification of BrdU+ cells as a percentage of the total number of cells in three areas (A = blue, B = red, C = green) of the blastema in *Acomys* or the scar in *Mus* from D5 to D30. F-H) Representative images of D10 (F) and D25 (H) blastema tissue with EdU+, BrdU+ and double positive cells. Red arrows = EdU+ cells, light blue arrows = BrdU+ cells, and white arrows = double positive cells (EdU+/BrdU+). G-I) Quantification of EdU+, BrdU+ and EdU+/BrdU+ cells as a percentage of the total number of cells in distal and proximal areas relative to the wound site at D10 (G) and D25 (I). n = 5 per time point per species. (* *p*<0.05).

We quantified healing tissue area from D10-30 which supported continued addition of new tissue during regeneration (Fig. 2B). In contrast, after a proliferative burst, no new fibrotic tissue was added in *Mus* ears after D10 (Fig. 2C).

Analyzing BrdU+ cells during regeneration revealed that from D15 to D30, the number of cycling cells was significantly higher in the distal areas compared to the proximal areas (injury site) while there was no difference between the distal-dorsal and distal-ventral areas (Fig. 2D and Supplementary Table 1). An exception was D20 when the new tissue areas were small or flat and thus distal and proximal areas were close (Fig. 2D). While cell proliferation in the proximal compartment decreased over time, proliferation in the distal compartment remained >10% from D10-30 contributing to the steady production of new tissue. (Fig. 2D). Quantifying BrdU+ cells in *Mus* ears, we observed a narrow extension of new tissue (Fig. 2A). After a burst of DNA replication (>25%) in *Mus* at D5, the percent of BrdU+ cells precipitously declined from D10-30 coinciding with no increase in tissue area over this time period (Fig. 2C, E).

To assess the spatial distribution of cycling cells during regeneration (D10-30), we first analyzed the epidermis where stem cell localization and proliferation are well known. Basal stem cells in the deeper layer of the epidermis show symmetrical and asymmetrical cell division depending on whether they produce progeny or more stem cells (Blanpain & Fuchs, 2006).

Analyzing our pulse-chase data, we observed stem, progenitor, and differentiating cells in the epidermis (Fig. S2A-B). The stratum basale contained slow-cycling stem cells (strong EdU+) and infrequently dividing stem cells (EdU+/BrdU+) while also indicating rapidly-cycling progenitor cells (strong BrdU+/diluted EdU+), and recently-cycling progenitor cells (strong BrdU+ Fig. S2A-B). In addition, we observed progenitor cells having acquired EdU that migrated apically withdrawing from the cell cycle and undergoing differentiation as per the absence of BrdU (Fig. S2B, yellow arrow). These cell behaviors demonstrated that our labeling scheme could effectively capture predicted stem cell behavior in a tissue with known proliferative properties.

Next we characterized proliferative behavior in the stromal compartment by quantifying cells that re-entered the cell cycle after injury but arrested within five days (EdU+ only) possibly to differentiate or re-enter the cell cycle later (slow-cycling progenitors), and cells that re-entered the cell cycle but maintained their proliferative state over the same period (EdU+/BrdU+ and BrdU+ only) in both distal and proximal areas relative to the wound site (Fig. 2F-I). We hypothesized that cell proliferation could either be clonal, with multiple daughter cells near proliferative progenitors in rosettes or could expand from the wound site. In the latter case, we predicted to capture progenitors cycling near the wound margin and daughter cells building up the blastema distally. At D10, we found a stochastic pattern of proliferation with a mixture of EdU+ cells, BrdU+ cells, and double-positive cells in distal and proximal compartments, where the EdU+ cell population represented the biggest population (>25%) (Fig. 2F-G). At D25, however, we were able to observe a pattern where rapid-cycling (BrdU+) and infrequently-dividing (double positive) cells represented half of the proliferative population (>9%) in the distal compartment with the other half (∼8%) being represented by potentially slow cycling progenitors (EdU+) (Fig. 2H-I). In comparison, the proximal area was represented mostly by EdU+ cells (5%) that may have started differentiation and a very small percentage of rapid-cycling and infrequently-dividing (1%) cells that could support expansion of some differentiating cell types (Fig. 2H-I). Supporting this pattern, we observed differentiation of new hair follicles and cartilage near the wound site. *In toto*, these results show that the proliferative progenitor population is maintained in a distal position relative to the wound site. Furthermore, our pulse chase labeling supports cell cycle progression and asynchronous cell proliferation during regeneration in *Acomys*, whereas thymidine analog incorporation without cell division was observed in *Mus*, suggesting an S-phase arrest after D10.

### The proliferative population comprises diverse cell types in Acomys

To determine cell state or identity among the proliferative population (PP), we co-labeled with cell type specific markers and EdU (see Methods). Although we found no mesenchymal cell types proliferating in uninjured tissue, we characterized the distribution of macrophages (IBA1+), chondroblasts (SOX9+) and two fibroblast subtypes (CRABP1+ or αSMA+) in comparison with *Mus* to identify any obvious differences in presence and location across tissue compartments (Fig. S3A-J). First, we found relatively few CD206+ and IBA1+ macrophages (Fig. S3A, B, F, G) nor αSMA+ cells which were sparse (Fig. S3E, J). We observed two populations of SOX9+ cells in both species: one associated with the perichondrium, and another associated with the sebaceous gland and hair follicle (Fig. S3C, H). Importantly, we observed a population of CRABP1+ fibroblasts scattered throughout the papillary dermis in *Acomy*s while these cells were not detectable in uninjured *Mus* skin (Fig. S3D, I).

Examining these same cell types at D5, we observed cell-cycle re-entry among many of them (Fig. 3A-J). By counting all the EdU+ cells in a counting frame and then quantifying double positive cells we estimated the fraction of cycling cells represented by each cell type. Because inflammation peaks at D5 (Gawriluk et al., 2020), we hypothesized that a large fraction of proliferating cells would be macrophages. Instead, IBA1+ and CD206+ cells represented only 5.3% and 4.95% of the PP respectively in *Acomys* and 5.3% and 0.96% in *Mus* (Fig 3A-D). In *Acomys*, the two fibroblast subtypes (CRABP1+ and αSMA+ cells) represented ∼14% and 18.65% respectively while chondroblasts (SOX9+ cells) represented ∼16% of the PP in *Acomys* (Fig. 3E, G, H). Surprisingly, although totally absent prior to injury in *Mus*, an emergent CRABP1+ population represented 28.9% of the PP at D5 (Fig. 3I). SOX9+ and αSMA+ cells represented 7.96%, and 33% of the PP in *Mus* (Fig. 3F, J). These data support that various cell types re-enter the cell cycle during regeneration *and* fibrotic repair.

**Figure 3.**
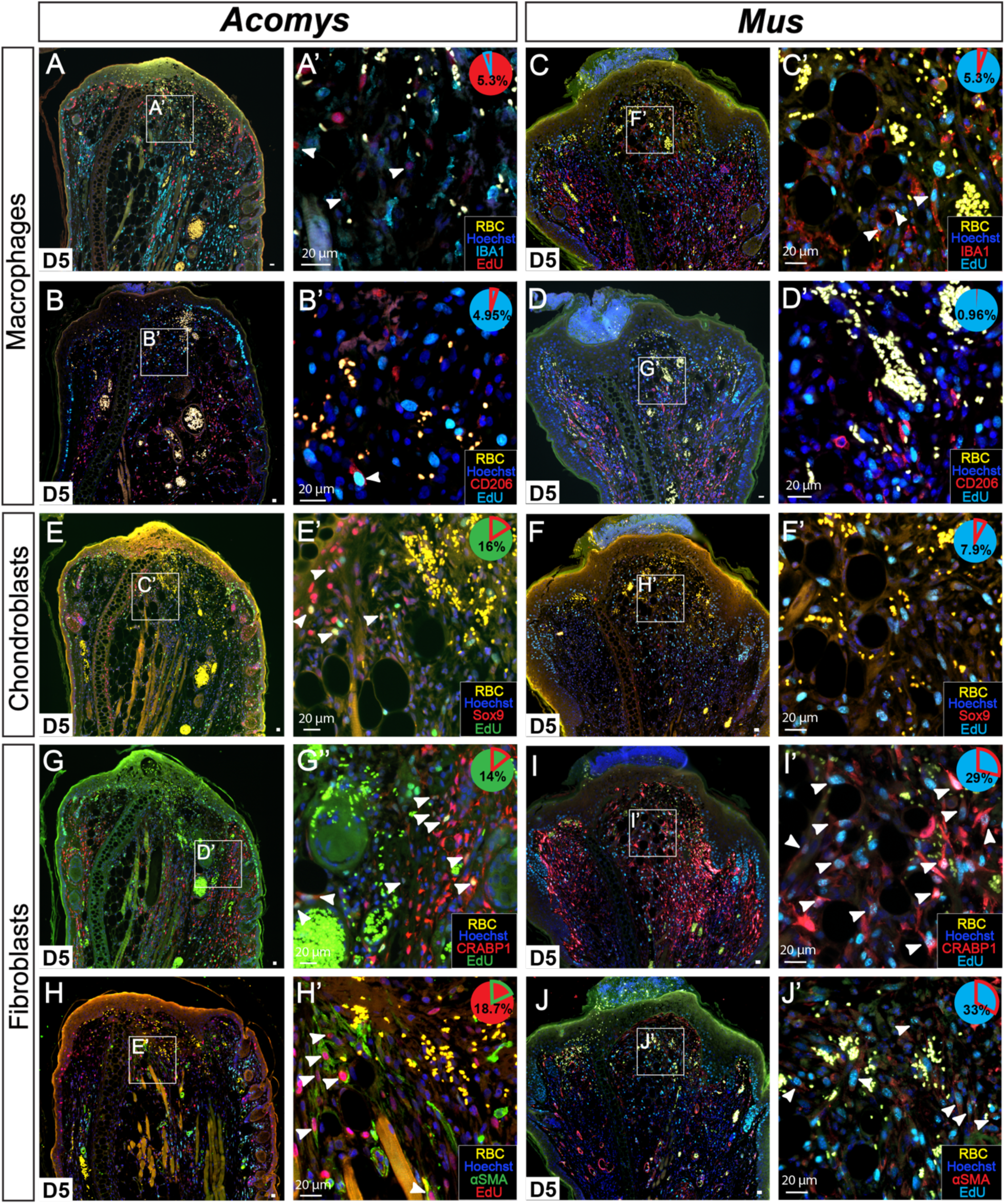
S-phase quantification of fibroblasts, chondroblasts and macrophages after injury at D5 in *Acomys* and *Mus*. A-D) Cycling macrophages double labeled with IBA1 and EdU in *Acomys* (A-A’) and *Mus* (C-C’) or with CD206 and EdU in *Acomys* (B-B’) and *Mus* (D-D’). E-F) Chondroblasts double labeled with Sox9 and EdU in *Acomys* (E-E’) and *Mus* (F-F’). G-J) Fibroblasts double labeled with CRABP1 and EdU in *Acomys* (G-G’) and *Mus* (I-I’). or with αSMA and EdU in *Acomys* (H-H’) and *Mus* (J-J’). White arrows indicate double positive cells (EdU+/cell type marker). Pie charts indicate percentage double positive cells of the total number of counted EdU+ cells. n = 5/species. RBC = red blood cells.

Only *Acomys* maintains a proliferative population during regeneration whereas activated cells did not appear to maintain cell cycle progression or appreciably divide during fibrotic repair. Co-staining with EdU, we confirmed the above-mentioned cell types remained part of the PP at D20 in *Acomys* (Fig. 4A-C and Fig. S4A-B). Of note, we observed that CRABP1+, αSMA+, and SOX9+ cells were segregated spatially throughout the blastema (Fig. 4A-C, G, H and Fig. S5A). While the CRABP1+ population was restricted to the papillary dermis at D0, this population expanded to deeper dermal areas at D5 (Fig. 3D, G and Fig. S5A) and occupied the dorsal blastema with another, smaller population in the ventral papillary zone at D20 (Fig. 4A-A’, G-H and Fig. S5A). In contrast, the αSMA+ fibroblast population appeared after injury in a distal and central position where reticular fibroblasts localize (Fig. 3H-H’ and Figs. S3E, S5A). This population shrunk at D20 and was largely non-overlapping with CRABP1+ cells (Fig. 4B-B’, G-H and Fig. S5A). Thus, as the blastema expanded, the αSMA+ fibroblasts maintained a small, distal population that disappeared in proximal tissue, while the CRABP1+ population expanded over time (Fig. 4G-H). In response to injury, the SOX9+ population that labeled the perichondrium appeared in the blastema perfectly aligned with the cartilage (Fig. 4C and Fig. S5A). IBA1+ and CD206+ cells were spread evenly across the blastema (Fig. S4A-B and Fig. S5A).

**Figure 4.**
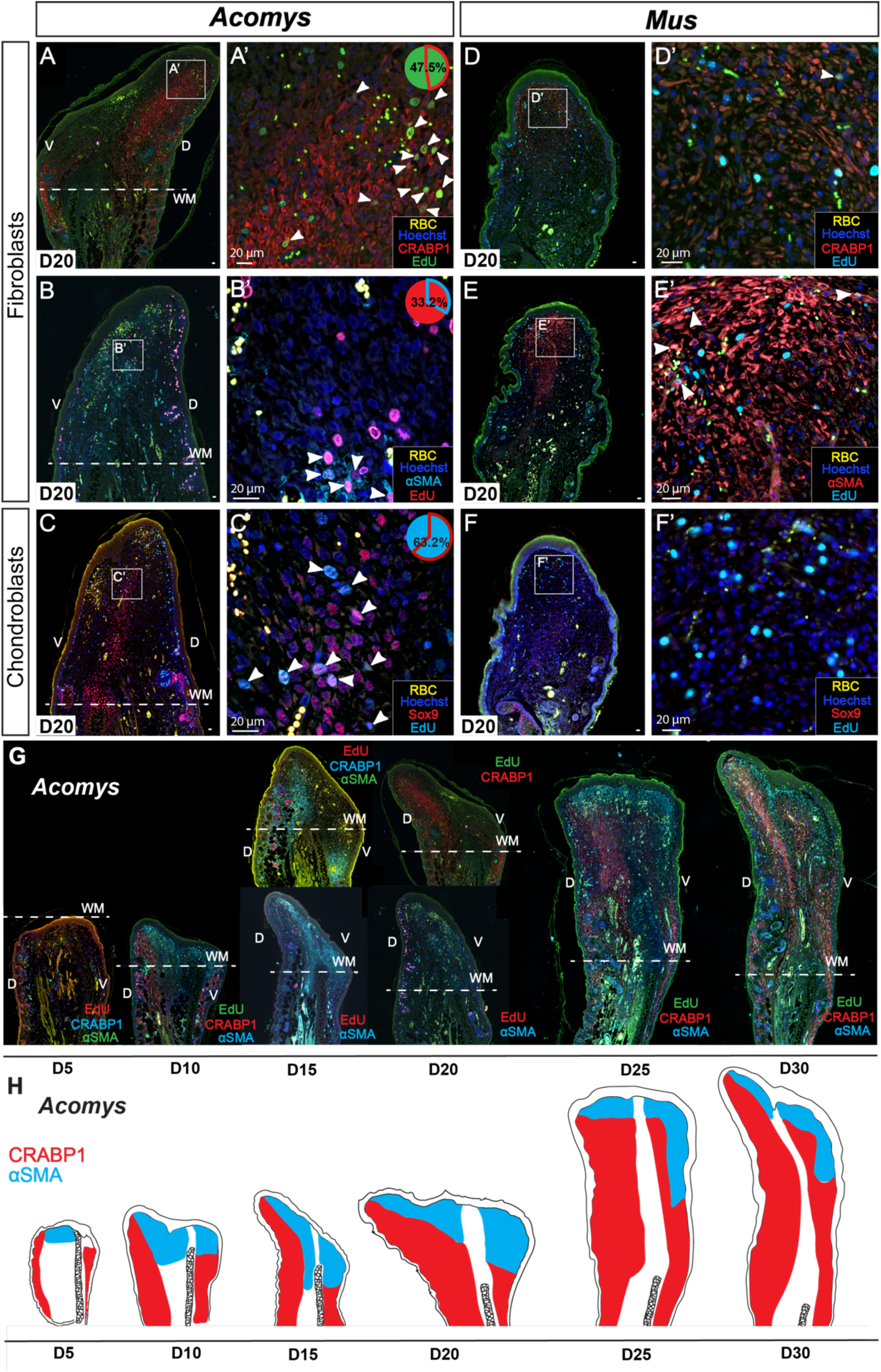
Spatial distribution of cell types within blastema or scar at D20. A-C) CRABP1+ (A-A’) or αSMA+ (B-B’) fibroblasts, and Sox9+ chondroblasts (C-C’) in *Acomys* at D20. D-F) CRABP1+ (D-D’) or αSMA+ (E-E’) fibroblasts, and Sox9+ chondroblasts (F-F’) in *Mus* at D20. G) CRABP1 and αSMA double staining in *Acomys* over time (D5-D30). H) Cartoon showing general distribution of CRABP1+ and αSMA+ fibroblasts in *Acomys.* White arrows indicate double positive cells (EdU+/cell type marker). Pie charts indicate percentage double positive of the total number of counted EdU+ cells. n = 5 per species. RBC = red blood cells.

The configuration of these populations was notably different in *Mus* during healing (Fig. 4D-F and Fig. S5B). Absent at D0, a CRABP1+ population emerged throughout the injury zone at D5, only to become restricted to the distal tip of the ear at D20 with distinctly fainter antibody reactivity (Figs. 3I & 4D and Fig. S5B). αSMA+ cells, absent in *Mus* at D0, became the dominant population in distal scar tissue (Fig. 4E and Fig. S5B). We observed some Sox9+ cells in the middle of the outgrowth (Fig. 4F and Fig. S5B), and IBA1+ and CD206+ cells distributed widely throughout the scar tissue (Fig. S4C, D and Fig. S5B). Western blotting for CRABP1 supported our observations using immunohistochemistry (Fig. S5C). Together, these results uncovered a distinct spatial segregation of cell types with a prominent CRABP1+ population present in *Acomys* before and after injury, that was merely transient in *Mus*.

### Species-specific cell states separate scaring and regeneration phenotypes

To extend our cellular studies, we next interrogated a scRNA-seq dataset harvested from uninjured (D0) and healing ear tissue at D3, 5, 10 and 15 from *Acomys* and *Mus* (Simkin et al., 2024). Unbiased clustering of batch corrected data for all time points (see Methods) revealed the same cell populations across species (Fig. S7). Thus, we extracted the four main stromal cell populations, fibroblasts (cluster 18), chondroblasts (cluster 19), lymphatic endothelial cells (cluster 16) and vascular endothelial cells (cluster 9), and generated a new UMAP (Fig. 5A). Re-clustering revealed eight subpopulations among the four original stromal populations: two populations each of vascular and lymphatic endothelial cells, three fibroblast subtypes, and a single chondroblast population (Fig. 5B). To investigate markers from our immunohistochemical analysis, we plotted *Crabp1*, *Acta2* (αSMA), and *Sox9* across cells on this UMAP (Fig. 5B).

**Figure 5.**
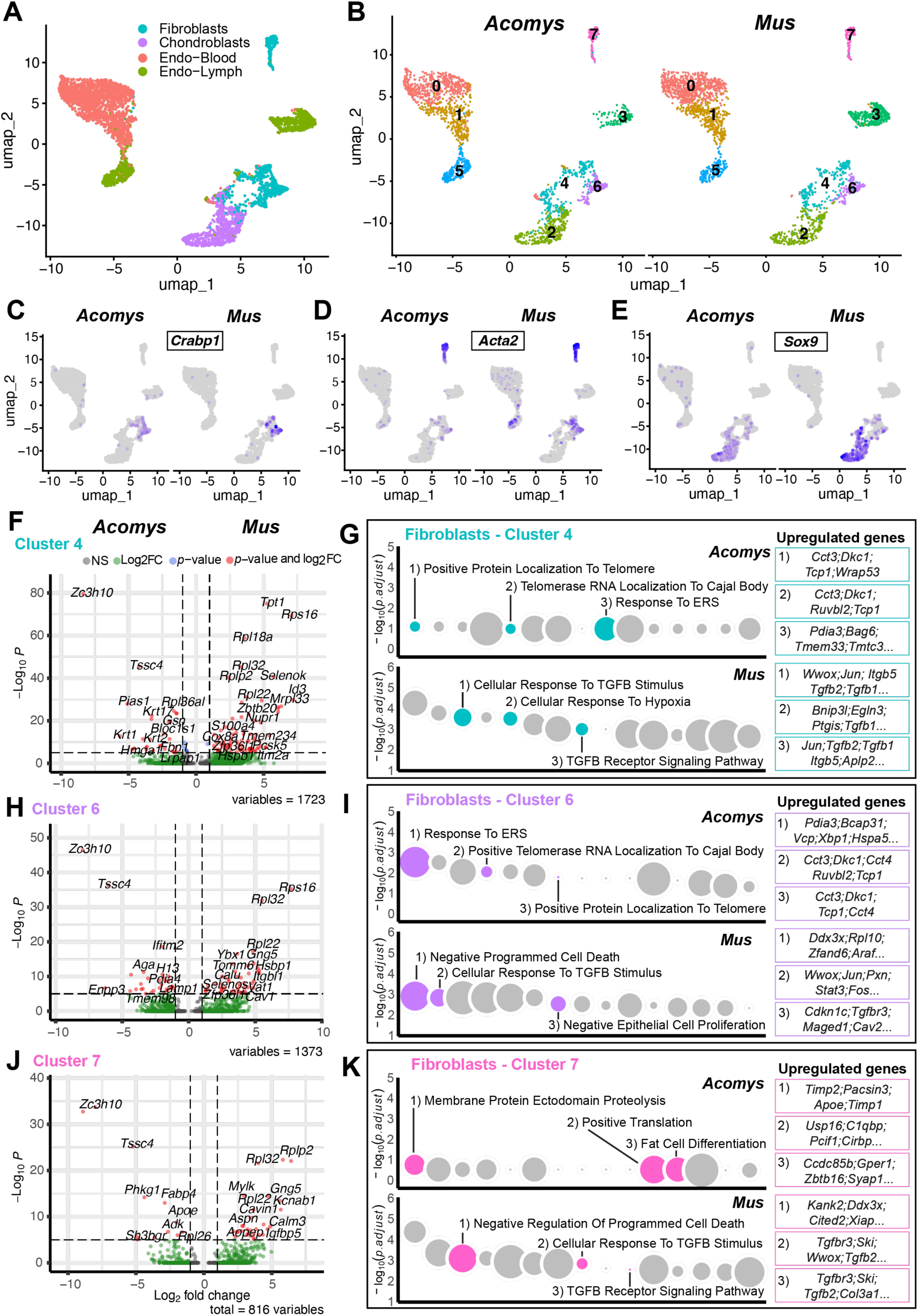
Single-cell analysis reveals different fibroblast cell states in *Acomys* and *Mus*. Cell suspensions collected from healing tissue preps representing D0, 3, 5, 10 and 15 were batch-corrected and plotted together (Fig. S7) (see Methods). A) UMAP projection of stromal cell types re-clustered from all cells (Fig. S7) from both species. B) UMAP projection of stromal subpopulations separated by species. Clusters represent subpopulations of major subtypes indicated in A. C-E) Feature plots for *Crabp1* (C), *Acta2* (D), and *Sox9* (E) separated by species. F, H, J) Volcano plots showing differentially upregulated genes between *Acomys* and *Mus* from clusters four (F), six (H), and seven (J) in B. Gene ontology Biological Process based on differentially upregulated genes between *Acomys* and *Mus* clusters four (G), six (I), and seven (K). Dot size represents the relative number of genes in each plot. Upregulated genes associated with each pathway are in color boxes. Refer to supplementary table S4 to see full list of genes.

Interestingly, we found that *Acta2* transcripts were strongly associated with one fibroblast subtype (cluster 7) while *Crabp1* was totally excluded from this cluster (Fig. 5B-D). *Sox9* marked chondroblasts in both species (Fig. 5E). To assess co-localization, we plotted *CRABP1* and *Acta2* together and found few double positive cells in *Acomys* (Fig. S8A). In contrast, fibroblast cluster six was mostly double-positive for *Crabp1* and *Acta2* in *Mus* (Fig. S8B). In addition, fibroblast cluster four and lymphatic endothelial cluster five were heavily enriched for *Acta2* in *Mus* (Fig. S8B). These data support a distinct *Acta2*+ fibroblast population in both species (cluster 7) and a *Crabp1+* fibroblast subtype in *Acomys* that is *Acta2-.* In *Mus*, *Crabp1* appears to only mark a transitional cell state that ultimately expresses *Acta2* while *Acta2* generally increases among fibroblasts and lymphatic endothelial cells supporting a fibrotic phenotype in *Mus*.

To better understand intrinsic differences between the fibroblast subpopulations, we ran differential gene expression (DGE) analysis comparing the three *Acomys* fibroblast clusters against the three *Mus* clusters (Fig. 5F-K). This analysis revealed 1497 genes (488 in *Acomys*, 1009 in *Mus*) differentially upregulated in cluster four, 1228 (510 in *Acomys*, 718 in *Mus*) in cluster six, and 743 (248 in *Acomys*, 495 in *Mus*) in cluster seven among both species (Fig 5F, H, J). Gene ontology of each cluster separated by species revealed a clear segregation in the fibroblast response to injury (Supplementary Table S4). The three fibroblast populations in *Mus* were associated with fibrotic pathways: response to TGFß stimulus, cellular response to hypoxia, and negative regulation of epithelial cell proliferation, among others (Fig 5G, I, K). In contrast, *Acomys* subpopulations were associated with telomere maintenance and cell proliferation pathways (Fig 5G, I, K). Interestingly, *Acta2*-expressing cluster seven was associated with regulation of fat cell differentiation, positive regulation of translation and regulation of membrane protein ectodomain proteolysis (Fig 5K). Chondroblasts (cluster two) followed a similar pattern with 2234 genes (786 in *Acomys*, 1448 in *Mus*) differentially upregulated. *Acomys* chondroblasts upregulated genes associated with translation, response to hydrogen peroxide and gene expression, while *Mus* chondroblasts upregulated genes associated with TGFß stimulus and regulation of TGFß-Receptor signaling (Fig. S9B). Of note, all four clusters (two, four, six and seven) in *Acomys* highly upregulated *Zc3h10* and *Tssc4* (Fig. 5F, H, J and fig. S9A). The first gene has been shown to boost mitochondrial function (Audano et al., 2018), while the second gene promotes cell survival and proliferation in breast cancer tumors (Chen et al., 2022).

Together, these data show that all four *Mus* subpopulations activate fibrosis-related pathways, while the four subpopulations in *Acomys* are biased toward pathways associated with cell proliferation and growth. Importantly, while the same markers may be able to identify fibroblast subtypes in both species, the intrinsic ability of each subtype to respond to injury differs.

### CRABP1+ fibroblasts do not induce regeneration in neonatal Mus

Previous studies using very large wounds in lab mice observed hair follicle regeneration in the wound center, an observation that has been attributed to the presence of *Crabp1+* fibroblasts (the so-called WIHN model) (Jiang & Rinkevich, 2021; Phan et al., 2021b). Since we did not observe CRABP1*+* cells in uninjured *Mus* tissue, but did observe them in *Acomys* ear dermis prior to injury and during regeneration, we hypothesized these cells disappear during *Mus* ear development. To test this hypothesis, we assessed CRABP1 reactivity in embryonic, neonatal, juvenile and adult ear tissue (Fig. 6A). In *Acomys,* we found CRABP1+ reactivity across all stages (embryonic and neonatal) and into adulthood (8 months old) (Fig. 6A).

**Figure 6.**
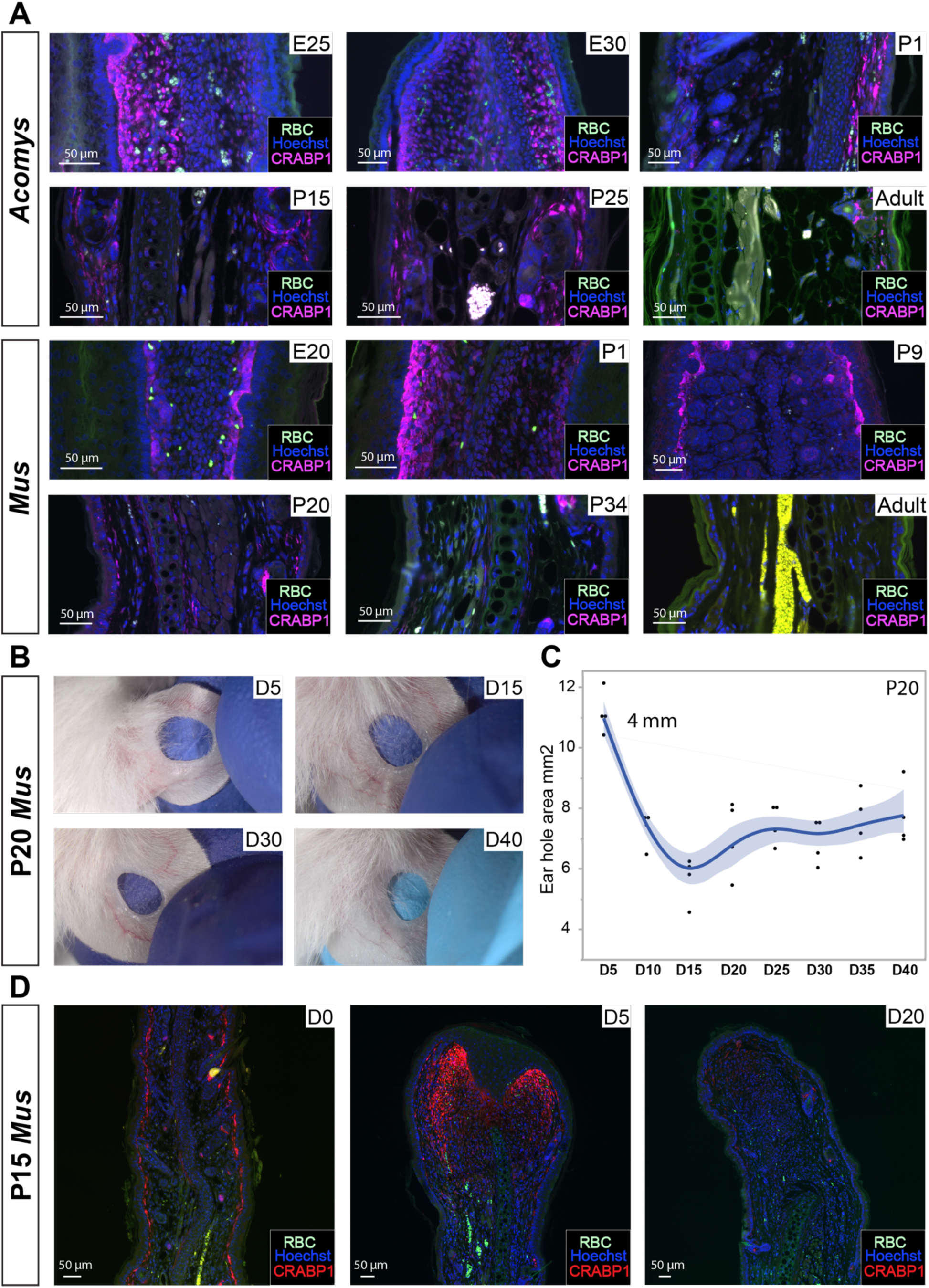
Tissue injury in neonatal mouse ears when CRABP1+ fibroblasts are present does not facilitate regeneration. A) Immunostaining for CRABP1 in *Acomys* and *Mus*. Age in days indicated in upper right corner of each image. B) Images showing representative ear hole closure from P20 injured *Mus* after a 4 mm ear punch. C) Quantification of ear hole closure area from D5 to D40. D) CRABP1 staining in P15 *Mus* before and after injury. n = 3 per species per time point. Green channel shows autofluorescence including red blood cells (RBC).

CRABP1 reactivity was strongest at embryonic stages and restricted to the dermis where it persisted into adulthood; mainly the papillary dermis (Fig. 6A). In contrast, we found *Mus* fibroblasts started to lose CRABP1 expression around P20 and it was totally absent from the ear pinna in three month old adults (Fig. 6A).

Based on these observations, we hypothesized that challenging *Mus* ear pinnae using our regeneration assay in the presence of CRABP1+ fibroblasts might enhance ear hole regeneration. Performing a 4 mm ear punch in P20 animals, we found neonatal ear holes produced tissue until D15 (Fig. 6B-C), but did not close. Similarly, we performed 2 mm punches in P23 animals and did not observe regeneration (Fig. S10A-B). Next, we injured P15 *Mus* which have CRABP1+ fibroblasts and collected tissue at D5 and D20. CRABP1 reactivity after injury was similar to that observed in adult *Mus* with increased expression at D5 which disappeared by D20 (Fig 6D). Again, we did not observe regeneration. Together, these data show the presence of a CRABP1+ fibroblast population is not sufficient to promote a regenerative response and instead supports species-intrinsic adoption of cells states that support regeneration.

## DISCUSSION

Here we conducted a comprehensive analysis of cell proliferation during regeneration and found cycling cells maintained a distal position relative to the original injury, while proximal cells exited the cell cycle and differentiated. These data revealed controlled regulation of the cell cycle in response to injury such that while amputation served as an activation trigger in both species, regenerating tissue maintained cell cycle progression while cells participating in fibrotic repair arrested. Importantly, proliferation in *Acomys* did not lead to tissue overgrowth demonstrating regulated shutdown of cell cycle progression. This unites *Acomys* with other highly regenerative vertebrates where tightly controlled cell proliferation is a hallmark of epimorphic regeneration in zebrafish caudal fins (Nechiporuk & Keating, 2002; Poleo et al., 2001) and newt limbs (Hays & Fischman, 1961; Wallace & Maden, 1976). Cell proliferation in injured mammalian organ stroma, apart from dedicated stem cell populations, is rarely observed except during tumorigenesis where cell proliferation is uncontrolled (Hanahan & Weinberg, 2011).

Although evolutionary explanations for the presence of regenerative ability among some vertebrate clades remains poorly supported, the presence of indeterminate growth has been offered as an underlying prerequisite (Froehlich et al., 2013; Sebens, 1987). This hypothesis claims that indeterminate growers maintain populations of somatic stem or progenitor cells that behave like slow-cycling stem cells to support growth. There are two arms to this hypothesis. First is the presence of cycling somatic cells during homeostasis lowers the barrier to cell cycle re-entry upon injury. Second is that once cycling, cells do not activate tumor suppressor machinery during S-phase progression to initiate cell cycle arrest. These ideas partly stem from culture experiments with newt myotubes that easily re-enter the cell cycle after serum stimulation in contrast to mouse myotubes formed from C2C12 cells which are refractory to cell cycle re-entry (Tanaka et al., 1997; Tanaka & Brockes, 1998). Although *Mus* myotubes can be forced to re-enter the cell cycle, this requires deletion or inactivation of tumor suppressors retinoblastoma protein (pRB) and p16^ink4a^ (Gu et al., 1993; Schneider et al., 1994).

To examine if cycling somatic cells exist in highly regenerative vertebrates, we analyzed cell cycle re-entry and progression prior to and after injury between determinate (*Acomys*, *Mus*) and indeterminate growers (*Ambystoma* and *Danio*). In line with previous observations (K. Johnson et al., 2018; Simkin & Seifert, 2018), we found a small population of slow-cycling stromal cells present during homeostasis in indeterminate growers while we failed to detect these cells in determinate growers. This cycling population, however, scarcely contributed to blastema formation in zebrafish. Instead, cells that re-entered the cell cycle in response to injury were the major contributors to regeneration. This result supports a previous study in zebrafish where slow-cycling cells marked for eight weeks before injury did not contribute to blastema formation (Nechiporuk & Keating, 2002). Importantly, our data show that cells in *Acomys* and *Mus* re-enter the cell cycle in response to injury independent of their regenerative ability. An open question is how cell cycle progression is controlled for almost forty days and then downregulated so as not to create tissue overgrowth. Considered *in toto*, our cross-species analysis rejects the notion that a determinate growth mode prevents cell cycle re-entry and supports that the contribution of slow-cycling cells to the blastema is minimal, at least in fish.

Our cross-species proliferation experiments also assessed whether cells on the other side of the animal were activated by injury (so-called systemic cell activation). Work in lab mice found that acute muscle injury elicits a systemic response in which contralateral-located stem cells (CSCs) re-enter the cell cycle and access a priming state named G_Alert_ (Rodgers et al., 2014). In contrast to quiescent stem cells (QSCs), CSCs in mouse show an increased propensity to cycle, although their numbers remain low in comparison with stem cells located at the injury site (Rodgers et al., 2014). A similar systemic response was observed in axolotl limbs contralateral to amputated limbs (Johnson et al., 2018). Using contralateral control tissue collected in our experiments, we looked for primed cells *in vivo* following injury. In contrast to muscle stem cells in *Mus*, we found a negligible stromal response in *Acomys* and *Mus* with less than 0.5% of contralateral ear pinna cells responding to a systemic signal. While we used immunofluorescence to quantify active or primed mesenchymal cells in tissue sections, previous studies in mouse muscle used FACS sorted satellite stem cells to assess *in vivo* BrdU incorporation. Supporting our observations in *Acomys* and *Mus*, we could not find a significant response in contralateral axolotl limbs or zebrafish pelvic fins, at least in the first 72hrs after injury. While these results are at odds with a single axolotl study, those data included epidermal cells in their cell counts and later time points (Johnson et al., 2018). Thus, our data do not support significant stromal cell cycle re-entry in uninjured contralateral tissue from the four species analyzed.

Contemporary vertebrate regeneration studies using transgenics have concluded that blastemas particular to an injured tissue are formed from a heterogeneous population of cells at the amputation plane (rev. in Seifert & Muneoka, 2018). Although various stem cell reservoirs have been discovered in different mammalian tissues, except for the digit tip, exploring cell type contributions during complex tissue regeneration in mammals has been impossible. While we could not identify stem cells with our methods, our data support that *Acomys* expands pre-existing cell types to participate in regeneration via cell proliferation. One exception in our analysis were αSMA+ fibroblasts which appeared after injury but did not maintain αSMA identity in newly differentiated tissue. Interestingly, our scRNA-seq data indicated that an *Acta2*-expressing population expressed genes related to the regulation of fat cell differentiation suggesting that some myofibroblasts may differentiate into adipocytes in a regenerative context. Similarly, αSMA+ myofibroblasts were shown to differentiate into adipocytes when exposed to bone morphogenetic protein (BMP) triggered by neogenic hair follicles in large skin wounds (Plikus et al., 2017). Further supporting our observations, recent data from another *Acomys* ear pinna regeneration study reported that αSMA fibroblasts disappear after wound closure with only ACTA2+ vascular smooth muscle cells remaining (Brewer et al., 2021). We too observed high expression of TGFβ pathway components in *Mus*, but not *Acomys* fibroblasts. Taken with the previously observed insensitivity of *Acomys* fibroblasts to TGFβ induced myofibroblast differentiation (Brewer et al., 2021), *Acomys* appears to support reduced myofibroblast activation. While these data suggest a possible differentiation trajectory for myofibroblasts, the exact dynamics of αSMA+ cells await further study in a regenerative context.

In contrast to their role during regeneration, the role of myofibroblasts during fibrotic skin repair is well known (rev. in Plikus et al., 2021). Previous studies in skin have described first wound responders as a large proportion of reticular fibroblasts including αSMA+ fibroblasts (Driskell et al., 2013) which rapidly seal the wound area promoting scaring and inhibiting hair follicle formation (Correa-Gallegos et al., 2019). These data support our observations in *Mus* where αSMA+ fibroblasts are the main cell population in newly generated scar tissue. Together, these data show that αSMA fibroblasts are dominant in scaring but seem to be transient in regeneration.

CRABP1+ papillary fibroblasts abundantly participate in the regeneration of neonatal skin and deer antler velvet and thus we interrogated their behavior in our ear pinna assay (Phan et al., 2021b; Sinha et al., 2022). In contrast to the ephemeral nature of the αSMA+ population in *Acomys*, we observed a stark difference in CRABP1+ cells during regeneration and fibrotic repair. While we were unable to detect CRABP1+ cells prior to injury in *Mus*, these cells appeared at D5 as a subset of cycling cells only to disappear by D20. In *Acomys*, however, we detected CRABP1+ cells prior to injury and this population expanded to participate in blastema formation. CRABP1 has been found in a variety of regeneration contexts. For instance, this protein is expressed in *Mus* papillary fibroblasts that expand in large, somewhat regenerative wounds (Guerrero-Juarez et al., 2019). Other examples are the regenerating wounds in newborn (P2) mice that have a higher proportion of CRABP1+ fibroblasts (Phan et al., 2021a) and the velvet skin on deer antlers which have regeneration-primed fibroblasts expressing genes putatively associated with a regenerative phenotype (e.g., *CRABP1, RUNX1,* and *PRSS35*) (Sinha et al., 2022).

Rather than the presence or lack of a specific cell type, our scRNA-seq data suggests that it may also be one of cell state. Although all four stromal subclusters express some of the same markers in *Acomys* and *Mus*, our DGE and GO analyses revealed that the four *Mus* subclusters express biological processes related to fibrosis. In contrast, all four subclusters in *Acomys* seem to use biological processes related with cell proliferation; high transcription and telomere maintenance and genes that boost mitochondrial function and cell survival. Specifically, a subpopulation in both species that highly expressed *Acta2* exhibited divergent transcriptomic responses. Future studies are necessary to address whether *Acomys* αSMA+ cells transiently become myofibroblast-like only to later differentiate into adipocytes as suggested by our GO analysis. Our finding that two subpopulations expressed high telomerase and endoplasmic reticulum stress (ERS) related activity supported the persistence of proliferation in these cells. Telomerase activity is low in most non-proliferating somatic cells in humans and mice, while high activity has been reported in highly proliferating immortal and cancer cells (Kim et al., 1994; Sha & Bacchetti, 1997). Moreover, inhibition of telomerase results in death of tumor cells *in vitro* and *in vivo* showing the importance of this enzyme for tumor growth (Hahn et al., 1999). Similarly, various ERS responses have been shown to promote cell proliferation and survival in various types of cancer (Zhang et al., 2024). These data suggest that *Acomys* can access and use pathways other adult mammals do not outside of tumorigenesis.

As a functional test of cell type versus cell state as a driver of regeneration, we assessed the role of CRABP1+ fibroblasts after injury in neonatal mice before this population disappears. Previous studies have shown that papillary CRABP1+ fibroblasts are enriched in regenerative neonatal human and mouse skin but are lost early during postnatal development (Mine et al., 2008; Sinha et al., 2022). Examining neonatal and early postnatal stages, we found CRABP1+ fibroblasts in the *Mus* ear pinna which persisted until approximately P20. Using our ear punch assay when CRABP1+ fibroblasts are present in the *Mus* ear pinna, did not improve ear hole closure or stimulate regeneration. These results support that the presence of a cell population alone may be less important that the ability for cells to acquire a specific cell state needed to facilitate regeneration. A recent examination of regeneration and scarring phenotypes across eleven murid rodents, underscored the binary nature of a regenerative phenotype in that species appear to either completely regenerate or not (i.e., no partial regeneration) (Riddell et al., 2025). This supports that complex tissue regeneration requires multiple features across stereotypical healing processes such as intrinsic cell cycle licensing (Gawriluk et al., 2016; Tomasso et al., 2023), goldilocks regulation of reactive oxygen species (ROS) (Saxena et al., 2019) and a regenerative skewed immune cell response (Gawriluk et al., 2020; Simkin et al., 2017, 2024).

Future work investigating specific cell states across more regenerative and non-regenerative mammals will undoubtably shed light on a blueprint for complex tissue regeneration in mammals.

## ACKNOWLEDGEMENTS

We would like to thank Josh Sarli, Brennan Riddell and Ava Musarra for animal husbandry. We thank all members of the Seifert lab for helpful discussions in developing the manuscript and Brennan Riddle for helping to create the GO graphs. This research was funded in part by grants from the NIH (R01AR070313) and Aligning Science Across Parkinson’s (ASAP-020495) through the Michael J. Fox Foundation for Parkinson’s Research (MJFF) to AWS. For the purpose of open access, the author has applied a CC BY public copyright license to all Author Accepted Manuscripts arising from this submission.

## AUTHOR CONTRIBUTIONS

EOR, RA, and AWS designed research. EOR, MA and RA performed research. EOR, MA and AWS analyzed data. EOR and AWS wrote the paper, and all authors commented on and edited the paper.

## AVAILABILITY STATEMENT

The data, code, protocols, and key lab materials used and generated in this study are listed in a Key Resource Table alongside their persistent identifiers (Supplementary Tables).

