## Supplemental figures for "Specific cell states underlie complex tissue regeneration in spiny mice"

### SUPPLEMENTARY FIGURES

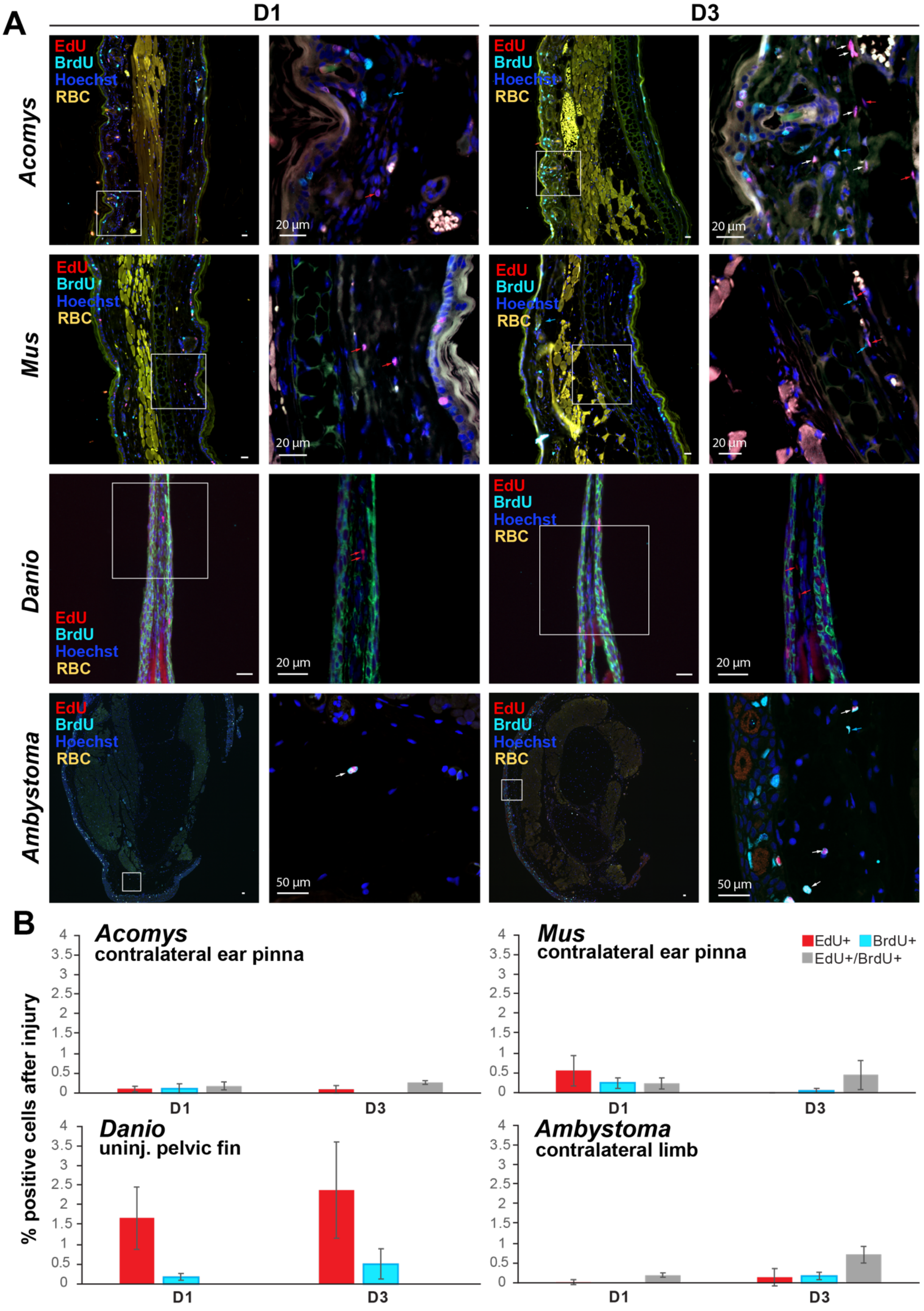
A
D1
D3
Acomys
Mus
Danio
Ambystoma
EdU
BrdU
Hoechst
RBC
20 µm
50 µm
B
% positive cells after injury
Acomys
contralateral ear pinna
Mus
contralateral ear pinna
Danio
uninj. pelvic fin
Ambystoma
contralateral limb
EdU+
BrdU+
EdU+/BrdU+
0
0.5
1
1.5
2
2.5
3
3.5
4
D1
D3

**Figure S1. Uninjured, contralateral tissue collected at D1 and D3 in *Acomys*, *Mus*, *Danio* and *Ambystoma.*** A) Representative images of uninjured, contralateral ear pinna tissue from *Acomys* and *Mus*, pelvic fin tissue from *Danio* and limb tissue from *Ambystoma.* Red arrows point to EdU+ cells, light blue arrows point to BrdU+ cells, and white arrows point to double positive cells (EdU+/BrdU+). RBC = red blood cells. B) Quantification for EdU+, Brdu+ and EdU+/BrdU+ cells as a percentage of the total number of cells counted at D1 and D3 post-injury in the four species. Images are a representation from n = 3 biological samples at each time point in *Acomys* and *Mus*. In zebrafish, n = 5 at D1 and n = 4 at D3. In axolotl, n = 6 at D1, n = 6 at D3.

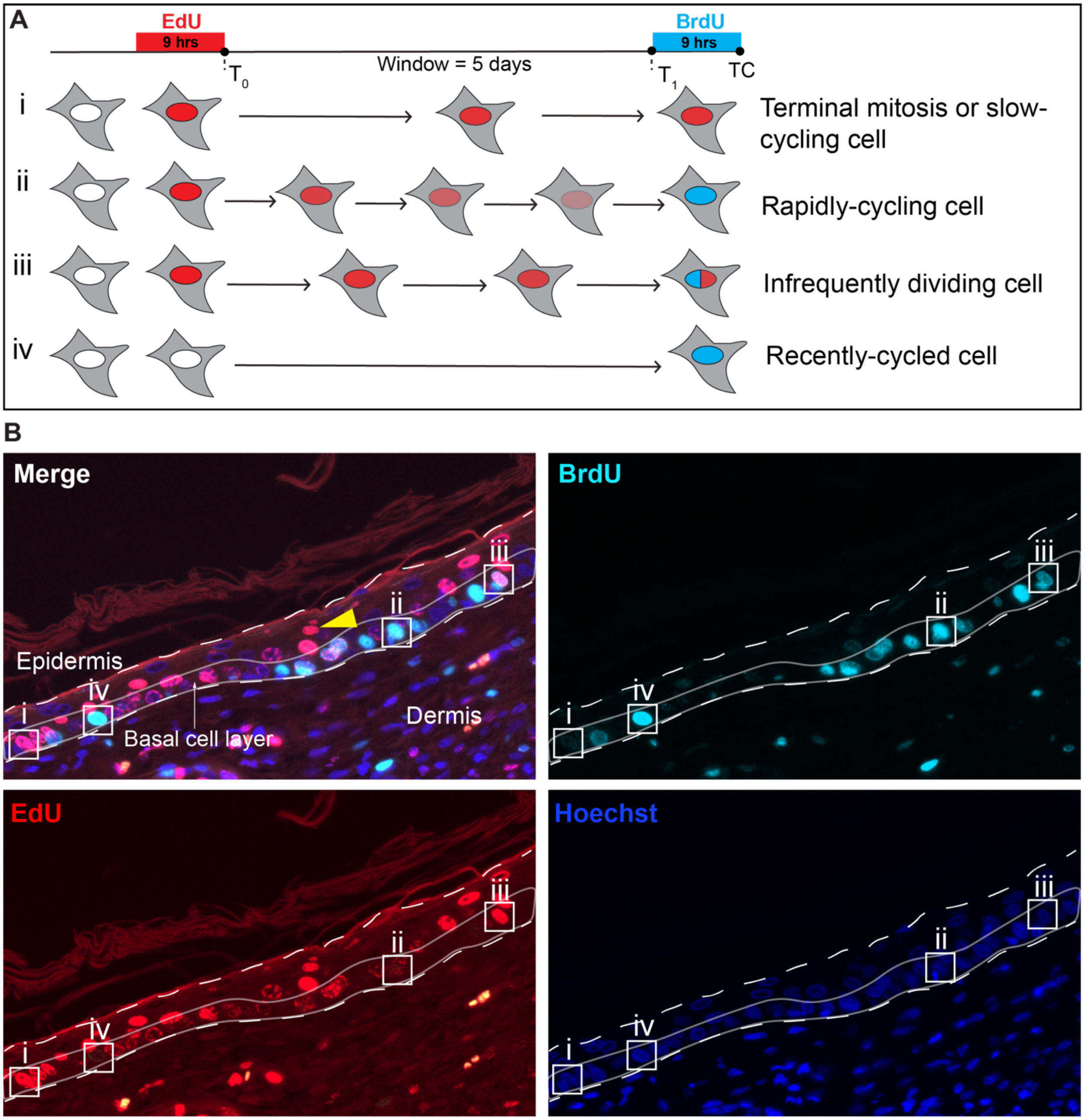


**Figure S2. Efficacy of EdU-BrdU pulse-chase method in epidermal compartment.** A) Scheme explaining pulse chase timing and possible labeling outcomes for cycling cells. B) Epidermal compartment from *Acomys* showing label incorporation in basal stem cells and differentiating progeny. Yellow arrow indicates EdU+ daughter cells outside the basal stem cell area depicting differentiation. i, ii, iii and iv in B refers to predicted labeling outcomes in A.

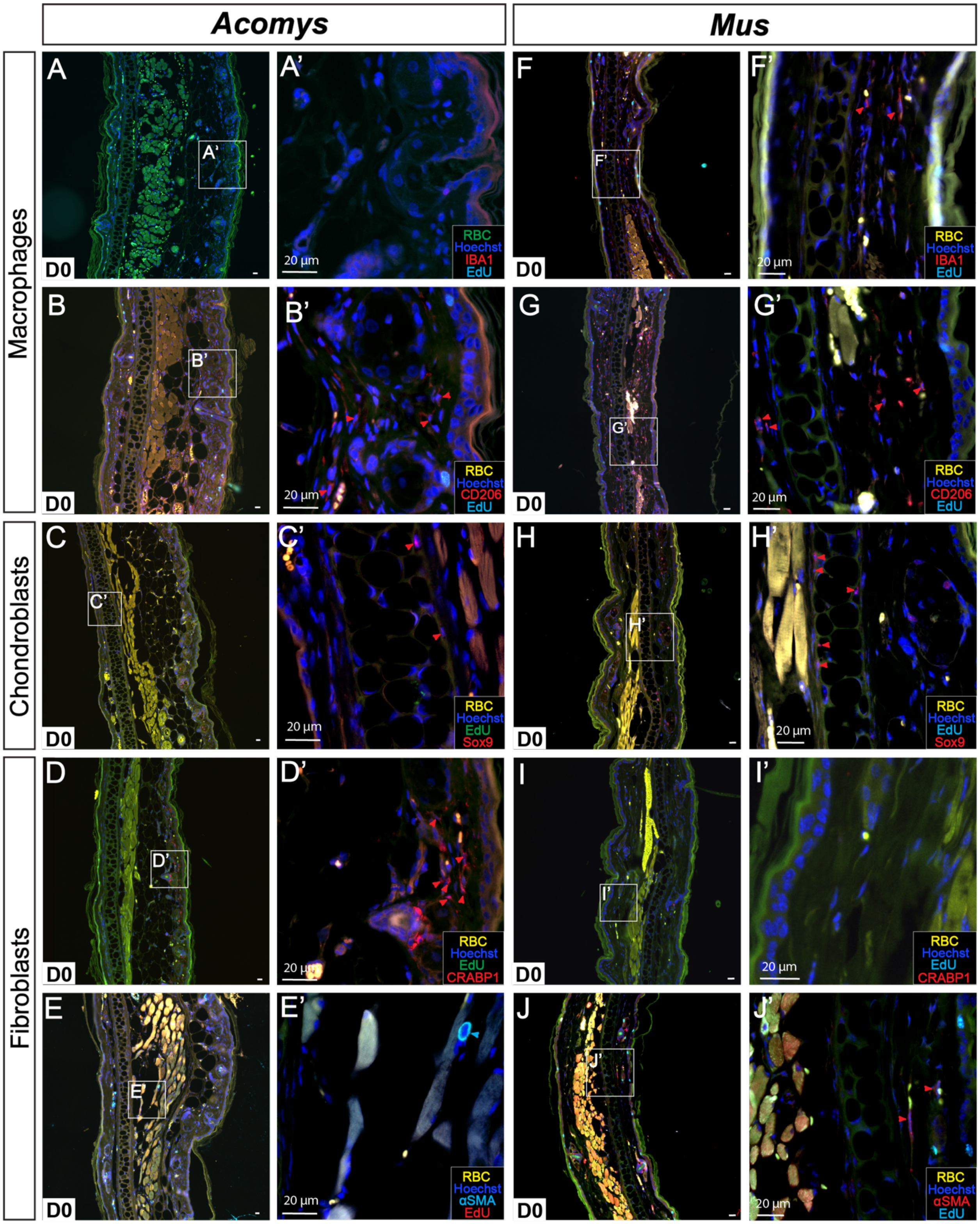


**Figure 3. Fibroblasts, chondroblasts and macrophages in uninjured tisse from *Acomys* and *Mus*.** A, F) Representative image of tissue double labeled with IBA1 and EdU in *Acomys* (A-A') and *Mus* (F-F'). B-G) Representative image of tissue double labeled with CD206 and EdU in *Acomys* (B-B') and *Mus* (G-G'). C-H) Representative image of tissue double labeled with Sox9

and EdU in *Acomys* (C-C’) and *Mus* (H-H’). D-I) Representative image of tissue double labeled with CRABP1 and EdU in *Acomys* (D-D’) and *Mus* (I-I’). E-J) Representative image of tissue double labeled with αSMA and EdU in *Acomys* (E-E’) and *Mus* (J-J’). Red and light blue arrows indicate positive cells for each marker. n = 5 per species. RBC = red blood cells.

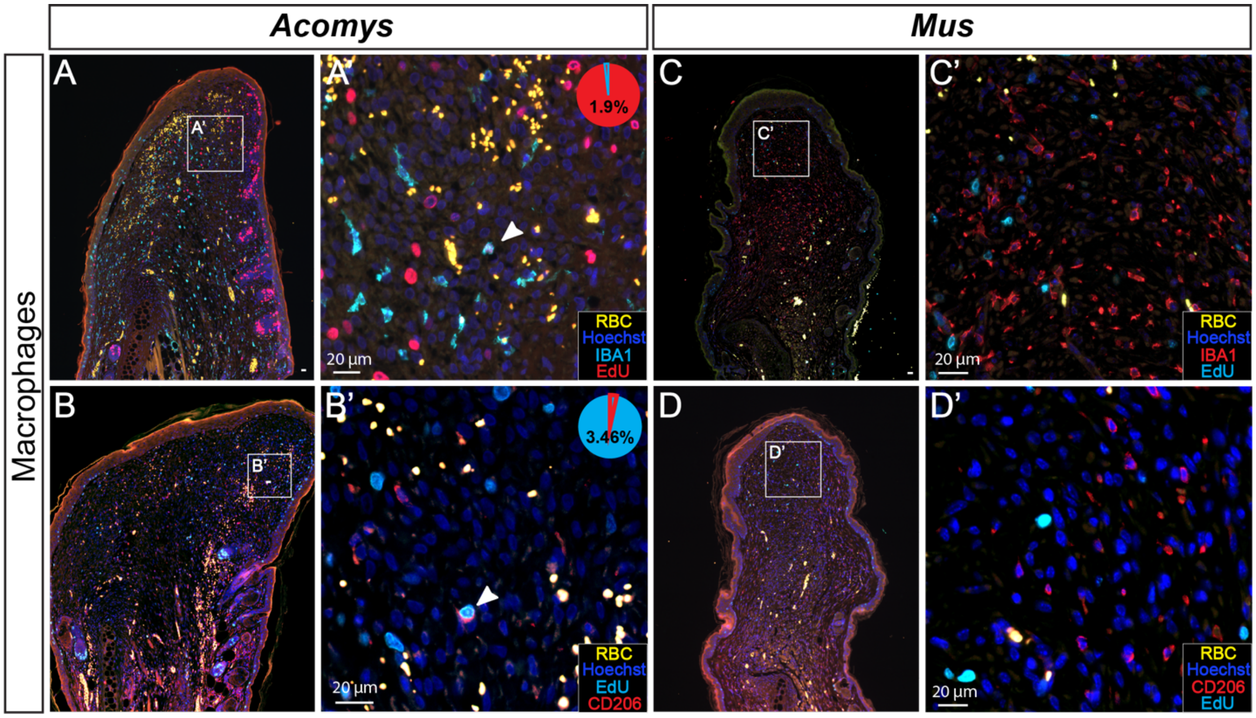


**Figure S4. Macrophages in blastema and scar tissue at D20.** A-B) Representative images of IBA1+ and CD206+ macrophages in the *Acomys* blastema. C-D) Representative images of IBA1+ and CD206+ macrophages in the *Mus* scar. White arrows indicate double positive cells (EdU+/cell type marker). Pie charts indicate the percentage double positive cells represent of the total number of EdU+ cells. n = 5 per species. RBC = red blood cells.

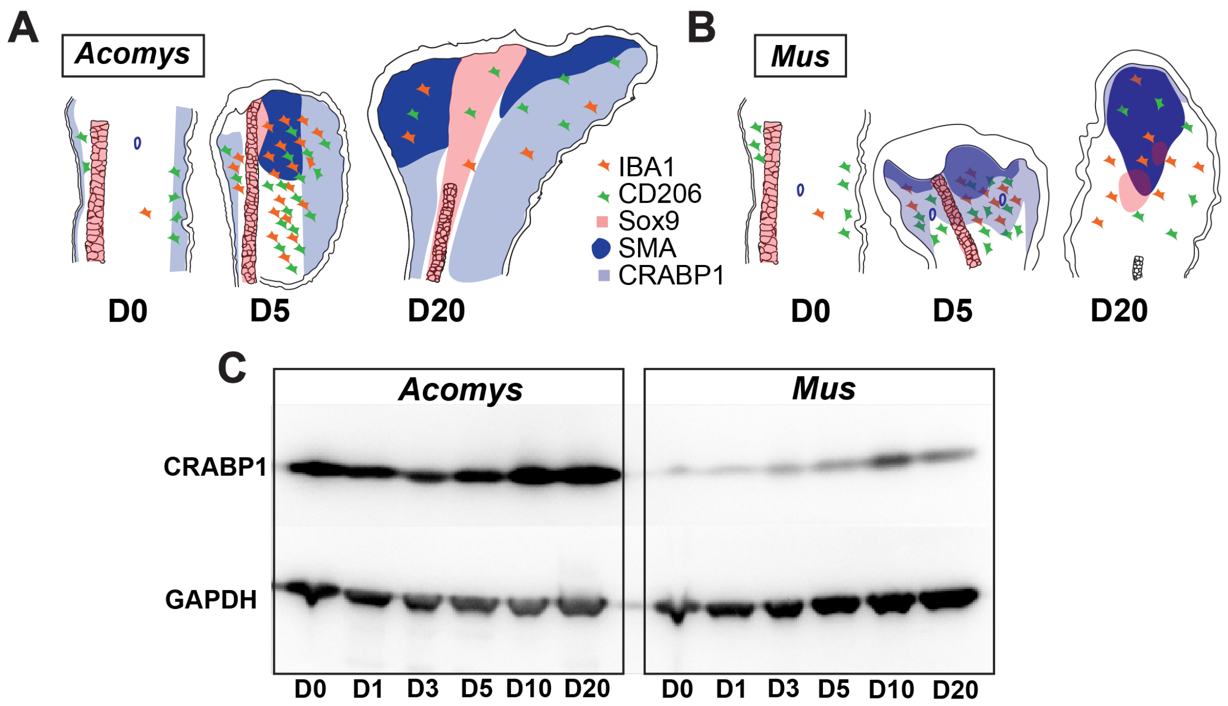


**Figure S5. Summary cartoon of cell type distribution in *Acomys* and *Mus* before and after injury.** A) Cartoon of cell type distribution at D0, D5, D20 in *Acomys.* B) Cartoon of cell type distribution at D0, D5, D20 in *Mus.* C) Representative western blot showing CRABP1 expression levels over time in *Acomys* and *Mus.*

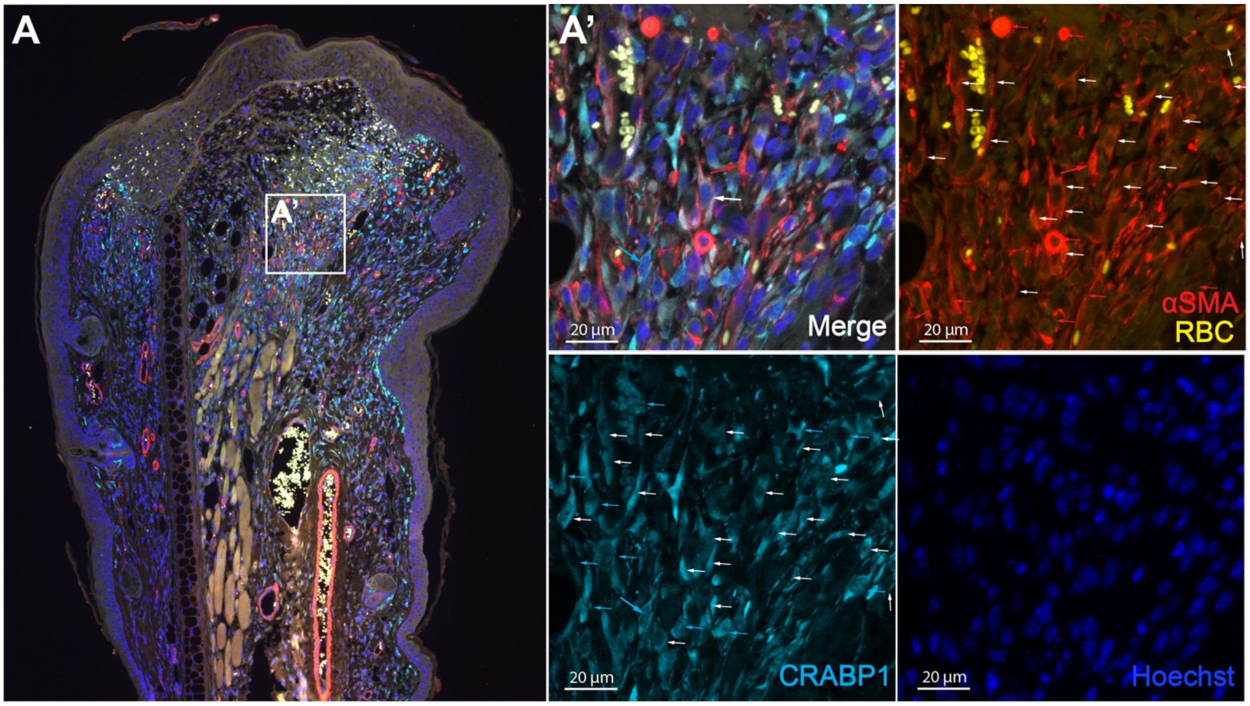


**Figure S6. CRABP1 and αSMA double staining in *Mus*.** A-A') Representative image of tissue double labeled with CRABP1 and αSMA in *Mus* at D5. Red arrows point to αSMA+ cells, light blue arrows point to CRABP1+ cells, and white arrows point to double positive cells (αSMA +/CRABP1+). n = 5 per species. RBC = red blood cells.

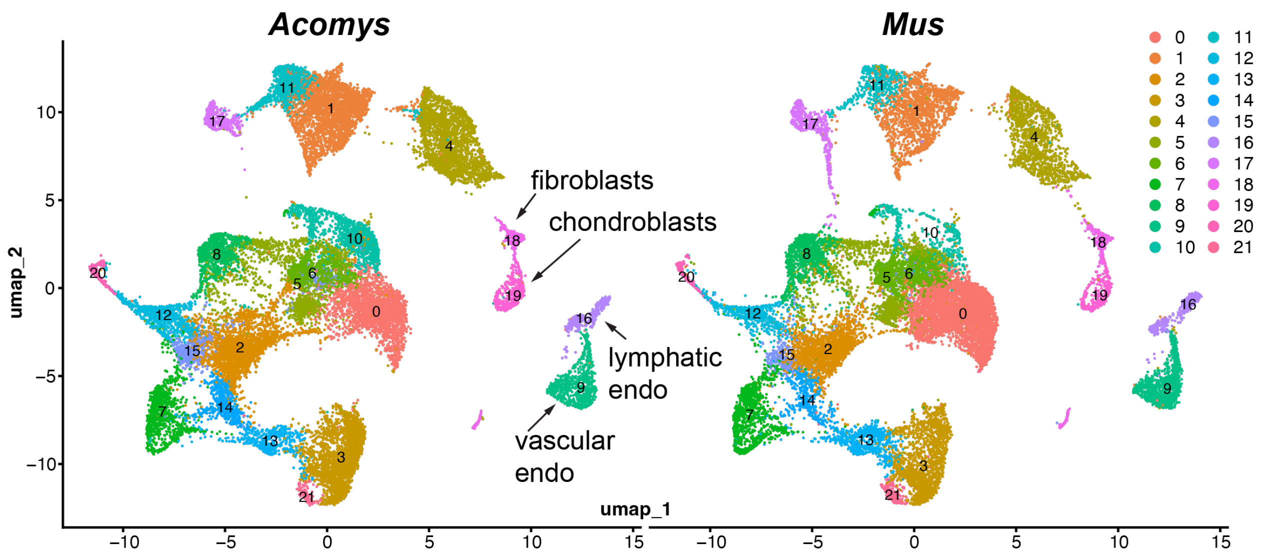


**Figure S7. Single-cell RNA-seq analysis of cells from healing tissue in Acomys and Mus collected at D0, 3, 5, 10 and 15 post injury.** A) Batch-corrected data and UMAP projection comprising all cell types in *Acomys* and *Mus* (see Methods for cell type identification). Major stromal cell types indicated on UMAP.

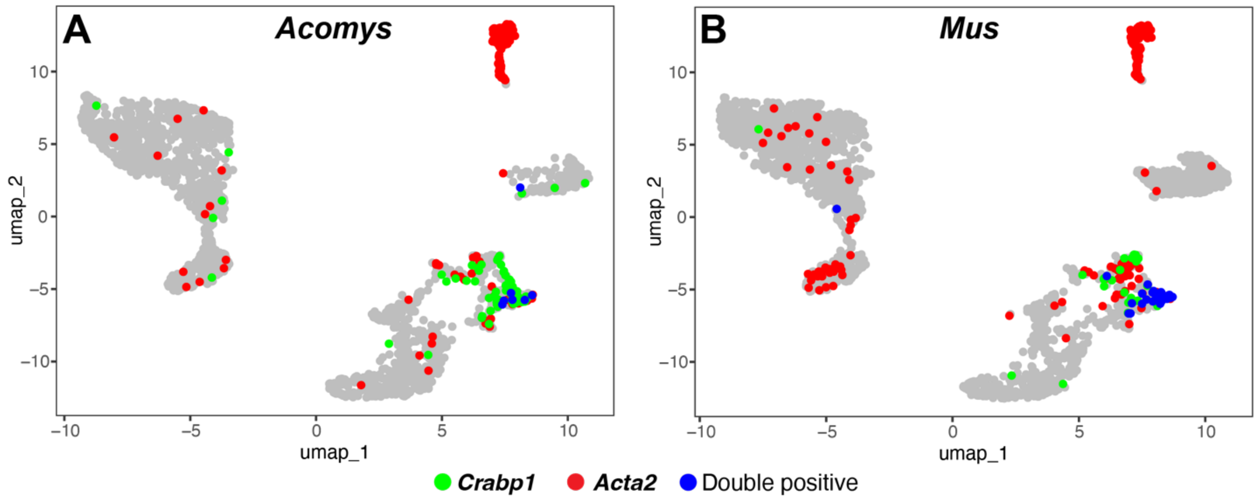


**Figure S8. Feature plots indicating single and double positive cells for *Crabp1* and *Acta2* in *Acomys* and *Mus*.** A-B) Feature plots of re-clustered stromal cells from Fig. S7 showing *Crabp1* (green), *Acta2* (red) and double positive cells (blue) in *Acomys* (A) and *Mus* (B).

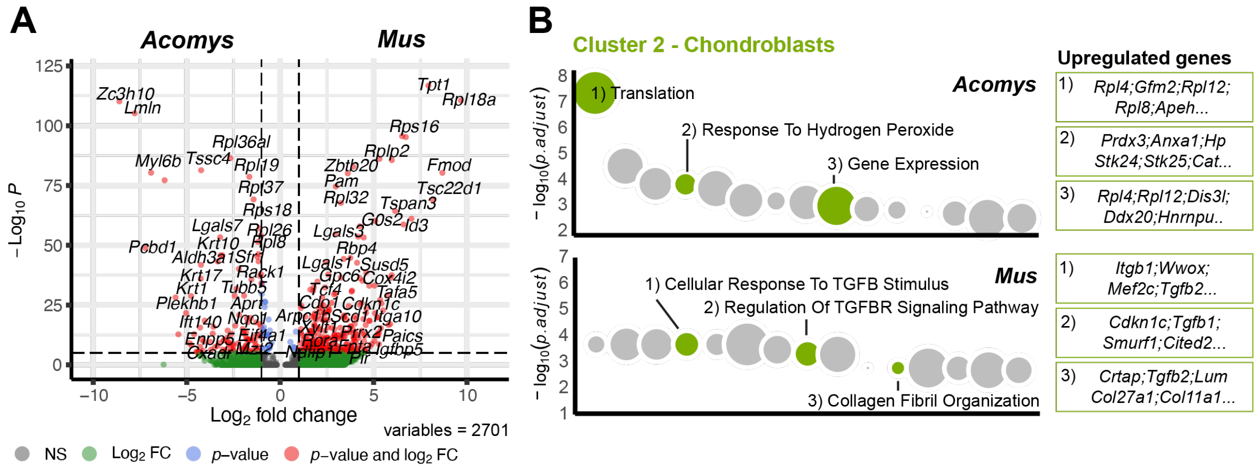


**Figure S9. Differential gene expression analysis reveals divergent cell states in *Acomys* and *Mus* chondroblasts.** A) Volcano plot of differently upregulated genes between *Acomys* and *Mus* chondroblasts. B) Gene ontology Biological Process based on differentially upregulated genes between *Acomys* and *Mus* chondroblasts (K). Dot size represents the relative number of genes in each plot. Upregulated genes associated with each pathway are in green color boxes. Refer to supplementary table S4 to see full list of genes.

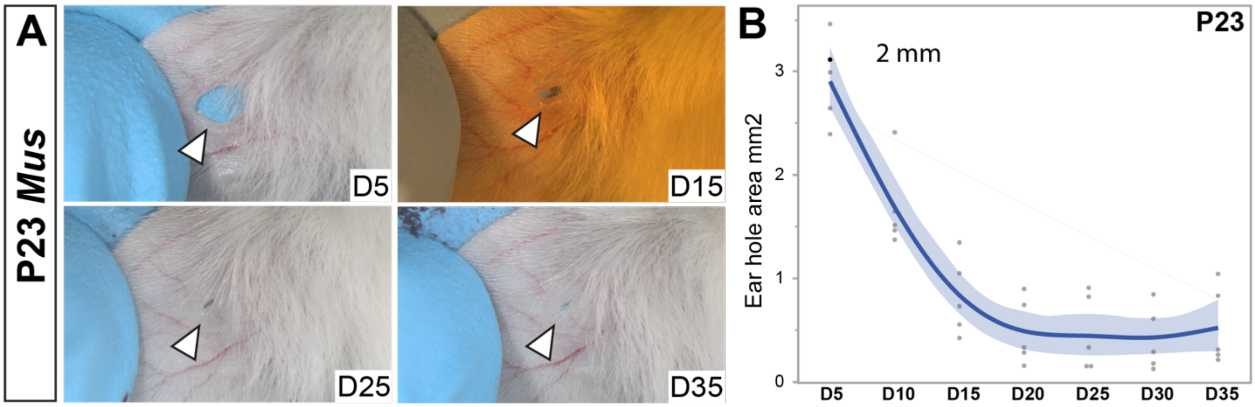


**Figure S10. Ear punch assay in P20 *Mus.*** A) Ear pinna healing over time after a 2 mm ear punch assay. B) Quantification of newly form scar tissue area after a 2 mm ear punch assay in P23 *Mus.* Each point represents an individual animal. n = 5 per time point per punch size.
